# Androgens Drive SOX9 Upregulation in Injured Proximal Tubular Cells

**DOI:** 10.1101/2025.10.23.684153

**Authors:** Corynne Vermillion Allison, Prisha S. Patel, Qiao Xuanyuan, Amanda Stayton, Josie A. Silvaroli, Isaac Z. Karel, Victoria C. Thorson, Gabriella Sloane, Helena Joseph Thailammanal, Yogesh Scindia, Christopher C Coss, Diana Zepeda-Orozco, Reena Rao, Subhashini Bolisetty, Sandeep K. Mallipattu, Benjamin D. Humphreys, Amandeep Bajwa, Navjot S. Pabla, Jiyoung Kim

## Abstract

**Introduction:** Sex influences susceptibility, recovery, and long-term outcomes after acute kidney injury (AKI) in humans. Rodent models have been invaluable for elucidating AKI mechanisms; however, most studies have focused on males, assuming direct applicability to females, an assumption that remains largely untested. In males, the transcription factor SOX9 regulates injury-associated proximal tubule cell states, supporting survival during injury but also contributing to maladaptive repair. Given the well-described role of SOX9 in sex determination during development and its emerging importance in AKI, we investigated whether it mediates sex-specific tubular responses in the adult kidney.

**Methods:** We first optimized differential ischemic and cisplatin conditions in males and females to generate injury-matched models. We then used these models to ask whether proximal tubular cell responses, particularly *Sox9* upregulation, are sex dependent. Tubule-specific knockout models of *Sox9*, *Sox4*, *Sox11*, *Vgf*, *Zfp24*, and the androgen receptor (*Ar*), along with gonadectomy and hormone replacement studies, combined with gene and protein analyses, were used to define regulatory networks.

**Results:** In both ischemic and nephrotoxic AKI, SOX9 expression was markedly blunted in females, with levels more than fivefold lower than in males. While injury biomarkers such as NGAL and KIM1 were equally induced in both sexes, *Sox9* and its downstream target *Vgf* showed markedly reduced induction in females, whereas the upstream regulator *Zfp24* was functionally relevant only in males. Deletion of *Sox9*, *Zfp24*, or *Vgf* worsened injury in males but not in females. In contrast, *Sox4* and *Sox11* were equally upregulated and protective in both sexes. Castration or proximal tubule-specific deletion of *Ar* in males abolished *Sox9* induction, establishing a testosterone-dependent regulatory axis.

**Conclusions:** These findings define a hormone-driven, male-specific tubular repair program and demonstrate that injury and recovery pathways differ fundamentally between sexes, underscoring the need for sex-inclusive therapeutic strategies for AKI.

**Translational Statement:** Sex influences susceptibility and outcomes after AKI, but the molecular basis remains unclear. SOX9, a transcription factor recently identified as one of the most highly upregulated genes in proximal tubular cells during AKI in both mice and humans, has been presumed to mediate a universal protective program. Using injury-matched murine models of AKI, we now demonstrate that this response is restricted to males, driven by testosterone and androgen receptor signaling, while SOX9 induction is markedly blunted and functionally dispensable in females. These findings reveal that tubular protective pathways differ fundamentally between sexes despite equivalent injury severity. Because SOX9 activation confers epithelial protection in males, therapeutic targeting of this pathway may have clinical relevance primarily in male patients. These data underscore the importance of incorporating biological sex into mechanistic, translational, and clinical studies of kidney injury and recovery.

## INTRODUCTION

Proximal tubular (PT) epithelial cells constitute nearly half of the kidney’s cellular mass and are the principal site of solute reabsorption and xenobiotic secretion.^1,2^ Their high metabolic demand renders them uniquely vulnerable to ischemia, toxins, and inflammation.^3^ PT injury is frequently the initiating event in the development of acute kidney injury (AKI), a common and multifactorial syndrome that primarily affects hospitalized patients.^4–7^ AKI is associated with substantial morbidity, mortality, and long-term progression to chronic kidney disease (CKD), yet remains without targeted therapies.^8,9^ Epidemiologic studies^10–13^ consistently reveal sex-based disparities, with males exhibiting greater susceptibility to AKI, although this appears to be etiology dependent. How proximal tubules mount injury responses in male versus female kidneys remains a critical, unresolved question with direct implications for therapeutic development.

Understanding the cellular and molecular responses of PT cells to injury is essential to define the mechanisms that drive injury, repair, and fibrosis. Rodent models have provided major insights into the pathogenesis of AKI, highlighting both protective and pathogenic pathways in epithelial,^6^ endothelial,^14–16^ and immune cells.^17,18^ However, most mechanistic studies have focused exclusively on male animals.^19^ Although some investigations^20^ have included both sexes and attributed differences to hormonal effects, they have largely overlooked a fundamental question: whether male and female PT cells activate similar transcriptional responses to equivalent injury.

The transcription factor SOX9 has emerged as a critical regulator of tubular injury responses.^21–23^ While largely absent under homeostatic conditions, *Sox9* is among the most highly upregulated genes in PT cells following AKI, as identified by single-cell and spatial transcriptomic studies in rodent models.^21–23^ We and others have shown that PT-specific deletion of *Sox9* does not affect baseline renal function but exacerbates injury in models of ischemia–reperfusion, cisplatin nephrotoxicity, and rhabdomyolysis.^21–25^ Conversely, delayed *Sox9* inhibition after the acute phase mitigates fibrosis and slows progression to CKD,^26^ suggesting a dual role for SOX9 in promoting survival during injury and driving maladaptive repair during chronic remodeling. Notably, all gene expression and functional studies to date have been conducted in male mice, leaving the role of SOX9 in female kidneys undefined.

Beyond its role in renal injury, SOX9 is also a pivotal transcriptional effector in sex determination.^27–30^ During gonadal development, SOX9 is activated downstream of SRY in XY embryos, driving testis formation.^31–33^ In both mice and humans, SOX9 is necessary and sufficient for male sex differentiation; its loss causes male-to-female sex reversal, while ectopic expression can induce testis development in XX embryos.^29,34–36^ This well-established role in male sex determination raises the possibility that SOX9 may also function in a sex-dependent manner in the adult kidney.

Here, we investigated whether SOX9 upregulation and function during AKI are sexually dimorphic. We found that SOX9 induction in PT cells is markedly attenuated in females, and PT-specific *Sox9* deletion has no effect on injury severity in female mice. In contrast, male PT cells exhibit robust SOX9 induction during AKI, which is dependent on testosterone and androgen receptor (AR) signaling. These findings reveal a striking sex-specific transcriptional program in proximal tubules and identify SOX9 as a male-specific mediator of epithelial protection during AKI.

## METHODS

### Mice strains and breeding

Mice were maintained in a temperature-regulated facility with a 12-hour light/dark cycle and provided with a standard diet and water ad libitum. All experimental procedures involving animals were conducted in accordance with the institutional animal use protocol approved by the Institutional Animal Care and Use Committee. C57BL/6J, Vgf-floxed, Sox9-floxed, Zfp24-floxed, and Ggt1-Cre transgenic mice (strains: 000664, 030571, 013106, 029023, and 012841) were sourced from Jackson Laboratories. Sox11-floxed and Sox4-floxed mice were procured from GemPharmatech (strains: T010043 and T013071). Additionally, mT/mG mice, which express a membrane-targeted, two-color fluorescent Cre-reporter allele, were obtained from Jackson Laboratories (strain: 007676). The floxed or mT/mG mice were crossed with Ggt1-Cre transgenic mice to generate models for conditional gene knockout or GFP-labeled renal tubular epithelial cells. In these transgenic mice, Cre recombinase expression is initiated in renal tubular epithelial cells at 1–2 weeks of age. For all mouse cohorts, offspring were ear-tagged and genotyped at 3 weeks of age. Genotyping was performed using standard PCR-based techniques, and kidney-specific gene deletion was confirmed through western blot analysis.

### Experimental models of acute kidney injury

All studies were conducted using age-matched male and female mice at 10–12 weeks of age. Littermate controls were utilized in all experiments involving conditional knockout mice. Investigators performing outcome assessments, measurements, and quantifications were blinded to the genotype or treatment status of the mice. For ischemia–reperfusion injury (IRI) experiments, mice were anesthetized with isoflurane and placed on a surgical platform, where their body temperature was continuously monitored. The skin was sterilized, and the kidneys were exposed for the procedure. Bilateral renal pedicles were clamped for 28–34 minutes, after which the clamps were removed to initiate reperfusion. The incision was sutured to close the muscle and skin. To mitigate fluid loss, 0.5 ml of warm sterile saline was administered intraperitoneally. In cisplatin nephrotoxicity experiments, cisplatin (15–25 mg/kg) was administered via intraperitoneal injection. Blood samples were collected on days 0–3 via submandibular vein bleeding or on days 1–3 via cardiac puncture following carbon-dioxide asphyxiation. Renal tissues were harvested and processed for histological, gene-expression, and protein analyses. The lipopolysaccharide-induced sepsis model was generated by anesthetizing mice with isoflurane (1.5–2 % for induction, 1–1.5 % for maintenance) and administering lipopolysaccharide (LPS; Cat. No. L2630, Sigma-Aldrich) via intraperitoneal injection.^37^ All animals were euthanized 24 hours post-injection, and blood and tissue samples were collected for analysis.

### Evaluation of AKI severity

Renal injury was evaluated using both serum biomarkers (blood urea nitrogen and creatinine levels) and histological analysis (hematoxylin and eosin staining). Blood samples were collected from mice at specific time intervals, and measurements of blood urea nitrogen and creatinine were performed using the QuantiChrom Urea Assay Kit (Bioassay Systems, DIUR-100) and the Creatinine Enzyme-Linked Assay Kit (Abcam, ab65340). For sepsis experiments, plasma creatinine was measured by LC–MS/MS at the UAB–UCSD O’Brien Center for Acute Kidney Injury Research. For histopathological analysis, kidney tissues were collected and embedded in paraffin at designated time points, both prior to and following AKI induction. Tissue sections (5 µm) were stained with hematoxylin and eosin according to standard procedures. A blinded histopathological evaluation was conducted by examining ten consecutive fields at 100× magnification from each kidney section, with a minimum of three mice per group. Tubular damage was quantified based on the proportion of tubules exhibiting dilation, epithelial flattening, cast formation, loss of the brush border, loss of nuclei, and basement-membrane denudation. The severity of tissue damage was classified according to the percentage of damaged tubules as follows: 0, no damage; 1, <25 %; 2, 25–50 %; 3, 50–75 %; and 4, >75 %, as previously described.^38–42^

### Transdermal Glomerular Filtration Rate (tGFR) Measurement

GFR was assessed using clearance of FITC-labeled sinistrin monitored via a transcutaneous detector (MediBeacon). Mice were anesthetized with isoflurane, and fur was shaved and depilated. After a 1–2 day acclimatization period, transdermal GFR sensors were affixed to the skin using a double-sided adhesive patch. A FITC-sinistrin solution (15 mg/mL in physiological saline) was administered intravenously (retro-orbital) at 75 mg/kg. Following injection, animals were returned to their cages, and GFR measurements were recorded over 1–2 hours. Elimination kinetics of FITC-sinistrin were analyzed to calculate GFR for each animal.

### Hormone add back experiments

For hormone add-back experiments, ten-to twelve-week-old C57BL/6J mice were gonadectomized and allowed to recover for 14 days to ensure stable depletion of endogenous sex hormones, as previously described ^43^. To evaluate acute hormonal regulation of injury responses, gonadectomized males received a single subcutaneous injection of testosterone propionate (2.5 mg/kg in 10 mL/kg sesame oil; Sigma-Aldrich) 48 hours before ischemia–reperfusion injury, and gonadectomized females received a single subcutaneous injection of 17β-estradiol-3-benzoate (5 µg per mouse in 0.1 mL sesame oil; Sigma-Aldrich) 48 hours before injury; vehicle controls received sesame oil alone. Both regimens were designed as single-dose physiological replacement paradigms consistent with established protocols in hormone-depleted mice.^43^

### Gene expression analysis

Total RNA (1 µg) from renal cortical tissues or isolated cells was reverse-transcribed using the RevertAid First Strand cDNA Synthesis Kit (Thermo Fisher Scientific). Quantitative reverse-transcription PCR (qRT-PCR) was conducted on a QuantStudio 7 Flex Real-Time PCR System (Thermo Fisher Scientific), employing SYBR Green Master Mix and pre-designed gene-specific primers (Sigma). Relative gene expression was calculated using the comparative CT (ΔΔCT) method, with β-actin serving as the reference gene. For in vivo gene-expression profiling of renal tubular epithelial cells, GFP-positive tubular epithelial cells were isolated from kidneys of reporter mice expressing membrane-bound EGFP via anti-GFP antibody-{Citation}mediated selection followed by magnetic-activated cell sorting (MACS) using columns from Miltenyi Biotec, as described in previous protocols.^24^

### Protein analysis

Whole-cell lysates from renal cortical tissues were prepared using a modified RIPA buffer consisting of 20 mM Tris-HCl (pH 7.5), 150 mM NaCl, 1 mM Na₂EDTA, 1 mM EGTA, 1 % Nonidet P-40, 2.5 mM sodium pyrophosphate, 1 mM β-glycerophosphate, protease and phosphatase inhibitors, and 1 % SDS. Protein samples were analyzed via Western blotting using Invitrogen Bis-Tris gradient midi-gels, followed by chemiluminescent detection using ECL reagent (Cell Signaling). For Zfp24 immunoprecipitation, cell lysates were prepared in a modified RIPA buffer containing 0.1 % SDS and 0.2 % β-maltoside. Immunoprecipitation was performed with a Zfp24 antibody, and precipitated complexes were analyzed by Western blotting using Invitrogen Bis-Tris gradient mini or midi-gels with ECL detection (Cell Signaling). Primary antibodies used for Western blotting included anti-β-actin (Santa Cruz Biotechnology, 47778), anti-NGAL (Santa Cruz Biotechnology, 50351), anti-SOX9 (Abcam, EPR14335-78), anti-cleaved caspase-3 (R&D Systems, MAB835), anti-ZFP24 (Abbexa, abx239672), and anti-phospho-linker (BioLegend, 685702), all diluted 1:1000. Secondary antibodies from Jackson ImmunoResearch were applied 1:2000. Densitometric quantification of the blots was carried out using ImageJ software, and expression levels of target proteins were normalized to β-actin in the corresponding samples.

### Statistical analysis

Data in all graphs are presented as mean ± SD. Statistical analyses were carried out using GraphPad Prism. p < 0.05 was considered statistically significant. To calculate statistical significance between two groups, a two-tailed unpaired Student’s t-test was performed. For comparisons among three or more groups, a one-way analysis of variance (ANOVA) followed by Tukey’s or Dunnett’s multiple-comparison test was used. No outliers were excluded, and all experiments were repeated at least three times.

## RESULTS

### SOX9 upregulation is suppressed in female kidneys during bilateral IRI-associated AKI

Female mice are known to exhibit resistance to ischemia-induced AKI, which has led previous studies to extend ischemic durations in females to achieve similar injury levels between sexes. In the current study, we optimized experimental conditions with varying clamp durations, identifying a protocol that induced comparable injury in both male and female C57BL/6J mice (10–12 weeks of age). Specifically, a 28-minute clamp time for males and a 34-minute clamp time for females resulted in equivalent injury levels (**Fig. 1a**). Blood urea nitrogen (BUN) and serum creatinine measurements from 0 to 48 hours indicated similar AKI severity in both sexes (**Fig. 1b–c**). Histological analysis of H&E-stained kidney tissue revealed equivalent tubular damage in males and females at 24 and 48 hours (**Fig. 1d**). At 24 hours, transdermal glomerular filtration rate (tGFR) measurements showed similar decline in both sexes (**Supplementary Fig. 1a**). Kidney cortical gene-expression analysis of injury biomarkers, namely neutrophil gelatinase-associated lipocalin (NGAL or *Lcn2*), kidney injury molecule-1 (KIM1 or *Havcr1*), and epidermal growth factor (EGF), showed comparable renal dysfunction in male and female mice (**Fig. 1e and Supplementary Fig. 1b–c**).

**Figure 1.**
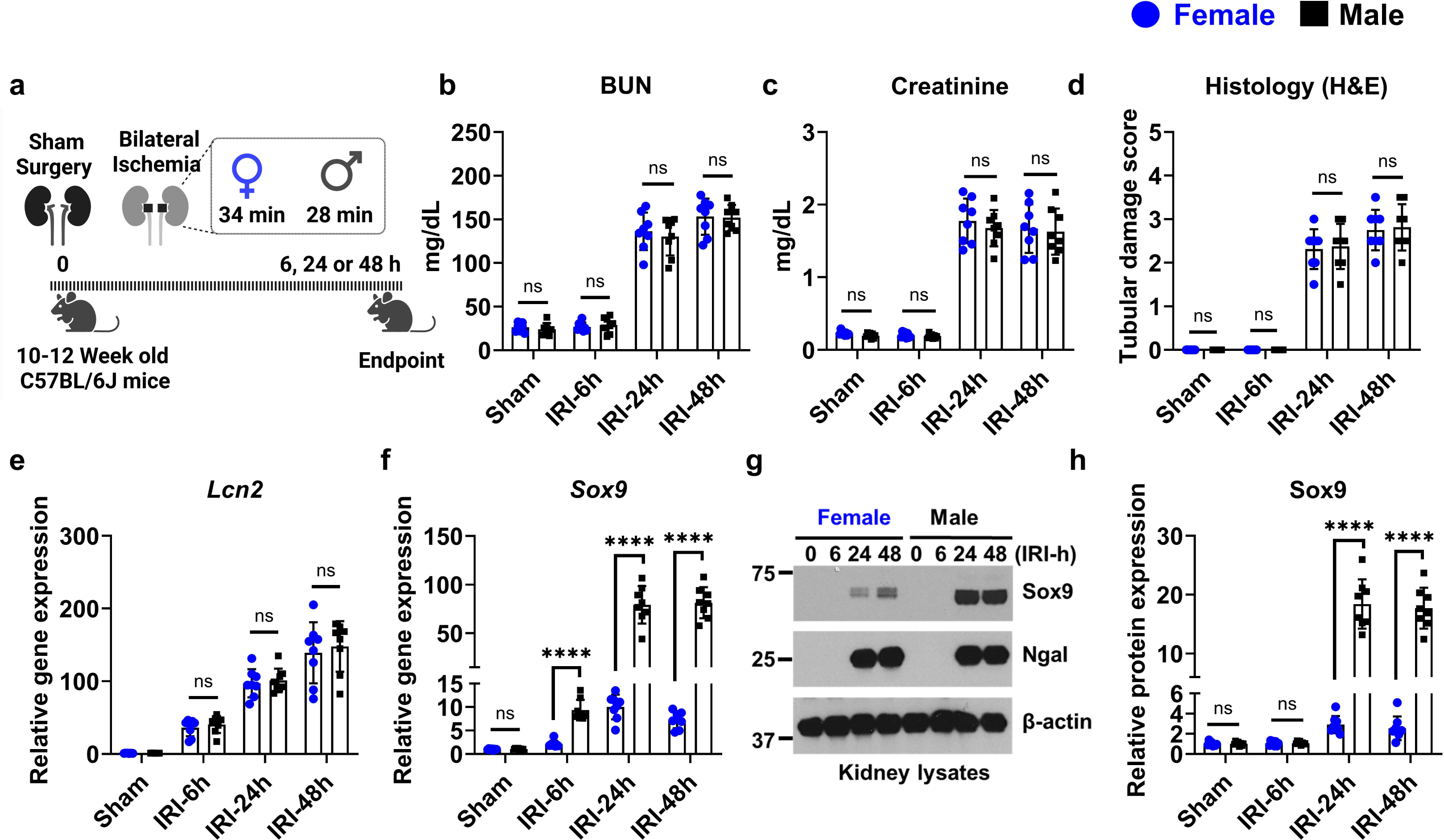
SOX9 induction is higher in males compared to females in injury-matched mouse models of bilateral ischemia-reperfusion. (**a**) Schematic of the experimental design. Age-matched (10–12-week-old) male and female C57BL/6J mice were subjected to bilateral renal ischemia (34 min for females and 28 min for males to produce injury-matched severity) or sham surgery. Serum and kidney tissues were collected at 6, 24, or 48 h post-injury. (b–d) Blood urea nitrogen (BUN) (**b**), serum creatinine (**c**), and histological injury scoring from H&E-stained sections (**d**) confirmed comparable degrees of kidney injury between sexes at all time points. (**e**) qPCR analysis of Lcn2 (NGAL), a marker of tubular injury, showed no significant difference between males and females. (**f**) Sox9 mRNA expression was significantly upregulated in both sexes following IRI but was markedly higher in males at 24 h and 48 h. (**g**) Immunoblot analysis of kidney lysates showing increased Sox9 and Ngal protein levels after IRI. Sox9 protein induction was greater in males. Ngal served as an injury control; β-actin was used as a loading control. (**h**) Densitometric quantification of Sox9 protein confirmed significantly higher expression in male mice at 24 h and 48 h post-injury. In all bar graphs (n = 6–8 biologically independent samples per group from 2–3 independent experiments), data are presented as mean ± S.D. Error bars represent 1 S.D. Statistical analysis was performed using two-way ANOVA followed by Tukey’s multiple-comparison test. *p < 0.05, **p < 0.01, ***p < 0.001, ****p < 0.0001; ns = not significant.

Once a model with equivalent AKI severity was established, we examined SOX9 expression at both the mRNA and protein levels in kidney cortical tissues. Quantitative PCR analysis revealed nearly tenfold lower induction of *Sox9* in female mice despite comparable injury between sexes (**Fig. 1f**). Immunoblot analysis further confirmed this blunted SOX9 upregulation in females, while NGAL was induced to similar levels in both sexes (**Fig. 1g–h**). Suppression of *Sox9* induction in female kidneys was also confirmed across multiple time points between 0–72 hours (**Supplementary Fig. 2**).

scRNAseq data analysis from prior studies ^44,45^ confirmed that proximal tubular SOX9 upregulation was blunted in female IRI-associated AKI (**Supplementary Fig. 3**). We also assessed SOX9 expression in models of cisplatin nephrotoxicity ^46^ and sepsis-associated AKI^37^, observing similar suppression of SOX9 upregulation in female mice under conditions where injury severity was comparable between sexes (**Supplementary Fig. 4**). Collectively, these results demonstrate that under equivalent injury conditions, SOX9 upregulation in proximal tubules is substantially blunted in female AKI.

### Proximal tubular SOX9 deletion increases male but not female susceptibility to IRI-AKI

We previously demonstrated ^24^ that deletion of *Sox9* in proximal tubules exacerbates AKI in male mice. To investigate its role in females, we generated PT-specific *Sox9* knockout mice by crossing *Sox9-floxed* mice with *Ggt1-Cre* mice. Because *Ggt1-Cre* is expressed postnatally,^47^ *Sox9* deletion does not cause renal abnormalities under baseline conditions. Baseline serum creatinine levels were unaffected by conditional *Sox9* deletion (**Supplementary Table 1**). After bilateral IRI, immunoblot analysis confirmed successful SOX9 deletion in both sexes (**Fig. 2a**).

**Figure 2.**
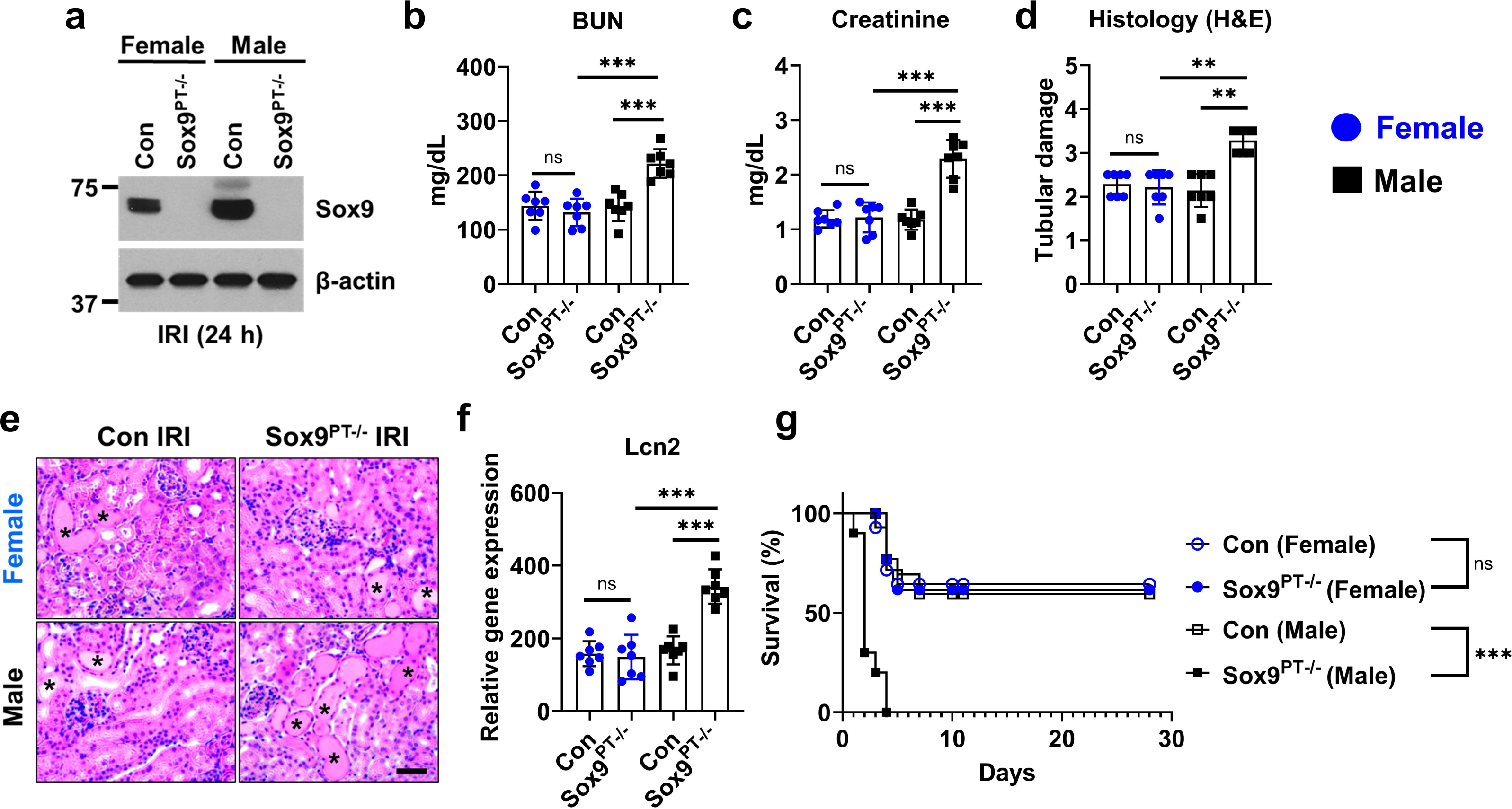
Tubular Sox9 deletion worsens ischemic kidney injury in males but not females. Sox9 conditional knockout mice (Sox9^PT-/-^) mice were generated by crossing Ggt1-Cre mice with Sox9-floxed mice to achieve tubular epithelial-specific Sox9 deletion. At 8-10 weeks age, littermate and Sox9^PT-/-^ were challenged with bilateral IRI to induce equivalent injury in control male and female mice. (**a**) Immunoblot of Sox9 in kidney lysates from control and Sox9^PT-/-^ male and female mice 24 h after bilateral ischemia-reperfusion injury (IRI), confirming Sox9 deletion. β-actin is the loading control. (**b–c**) Blood urea nitrogen (BUN) and serum creatinine measurements reveal significantly worsened kidney injury in Sox9^PT-/-^ male mice compared to controls, with no significant differences observed in females. (**d-e**) Histological tubular injury scores based on H&E staining showed exacerbated tubular damage in Sox9^PT-/-^ males. Representative H&E images depict damaged tubules with an asterisks. Scale bar: 100 µm. (**f**) qPCR analysis of Lcn2 (NGAL) gene expression confirms increased injury marker levels in Sox9^PT-/-^ males but not females. (**g**) Kaplan-Meier survival analysis shows significantly decreased survival in Sox9^PT-/-^ male mice following IRI, while females show no survival difference. All data are presented as mean ± S.D. with n = 6–8 biologically independent samples per group from three independent experiments. Statistical analysis was performed using two-way ANOVA with Tukey’s multiple-comparison test for panels (b–f), and log-rank (Mantel-Cox) test for survival (g). Significance is indicated by **p < 0.01, ***p < 0.001; ns = not significant.

In male *Sox9*^PT–/–^ mice, BUN and serum creatinine levels were significantly higher at 24 and 48 hours post-IRI, indicating more severe injury compared with controls (**Fig. 2b–c**). In contrast, no significant differences were observed in female *Sox9*^PT–/–^ mice (**Fig. 2b–c**). Histological analysis showed more extensive proximal tubular damage in male *Sox9*^PT–/–^ mice, while no differences were seen in females (**Fig. 2d–e**). NGAL (*Lcn2*) expression corroborated these findings (**Fig. 2f**). Long-term survival studies revealed increased mortality in male *Sox9*^PT–/–^ mice but no effect in females (**Fig. 2g**). Similar results were observed during cisplatin nephrotoxicity (**Supplementary Fig. 5**). These findings confirm that SOX9 protects male but not female kidneys during AKI.

### ZFP24 is the upstream regulator of SOX9 in males but not females

We recently identified ZFP24 as a key regulator of SOX9 upregulation in male proximal tubules. To determine whether ZFP24 also regulates SOX9 in females, we used PT-specific *Zfp24* knockout mice. Baseline BUN and creatinine levels were unaffected by *Zfp24* deletion (**Supplementary Table 1**). Successful gene deletion was confirmed by immunoblot analysis of kidney tissues (**Fig. 3a**). In male *Zfp24*^PT–/–^ mice, AKI severity was significantly exacerbated following bilateral IRI (**Fig. 3b–f**), confirming a protective role for ZFP24 in males. In contrast, no such effect was observed in female *Zfp24*^PT–/–^ mice, indicating that ZFP24 does not modulate kidney injury in females (**Fig. 3b–f**).

**Figure 3.**
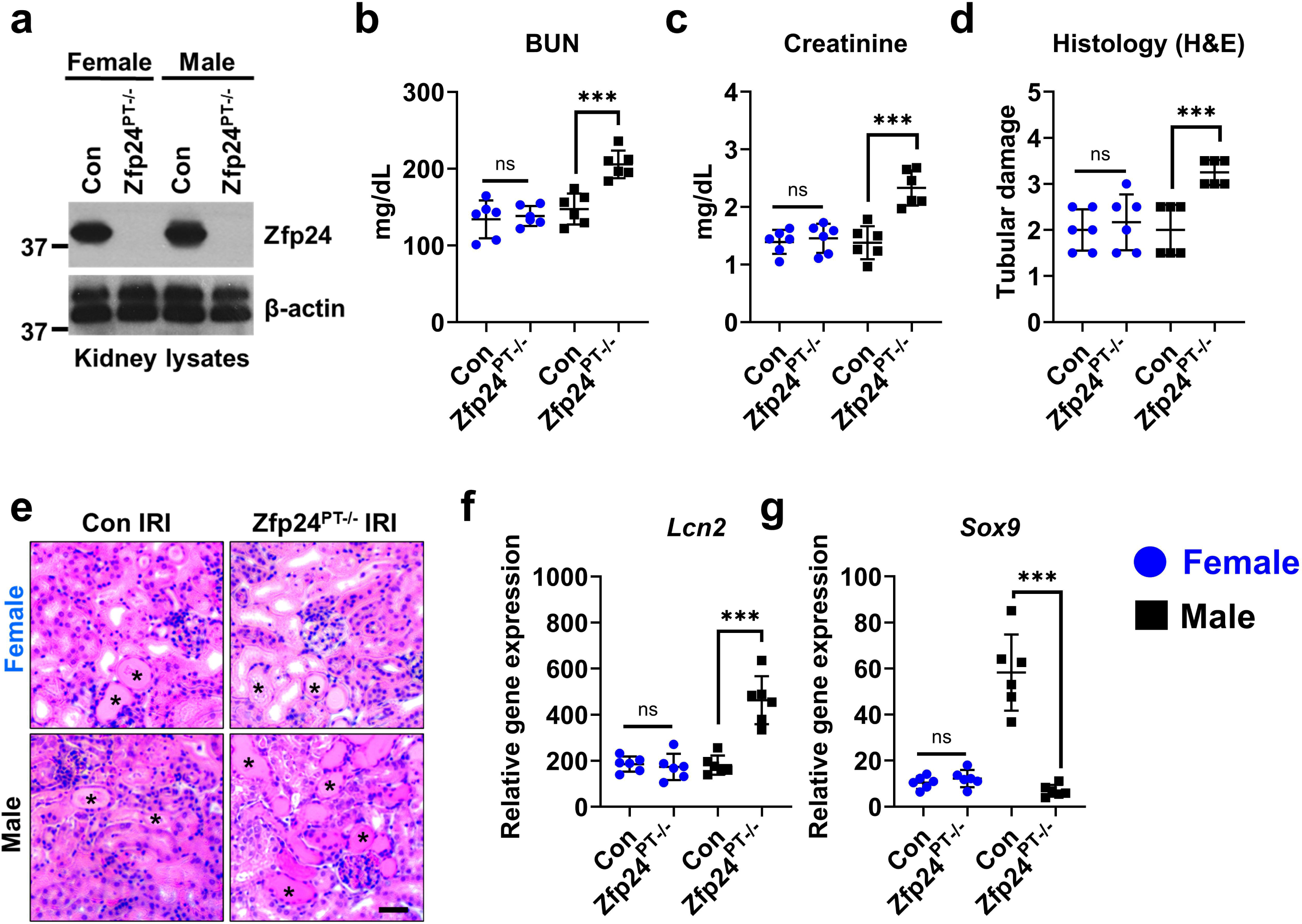
Tubular Zfp24 deletion worsens ischemic kidney injury in male but not female mice. Zfp24^PT-/-^ mice were generated by crossing Ggt1-Cre mice with Zfp24-floxed mice to achieve tubular epithelial-specific Zfp24 deletion. At 8-10 weeks age, littermate and Zfp24^PT-/-^ were challenged with bilateral IRI to induce equivalent injury in control male and female mice. (**a**) Zfp24 deletion was confirmed by immunoblot analysis of renal cortical lysates. (**b–c**) Blood urea nitrogen (BUN) and serum creatinine levels were significantly elevated in male Zfp24^PT-/-^ mice compared to controls, with no significant differences observed in females (n=6–9 per group). (**d-e**) Tubular damage assessed by histology (H&E staining) was increased in male Zfp24^PT-/-^ mice after IRI. Representative H&E images depict damaged tubules with an asterisks. Scale bar: 100 µm. (**f–g**) Relative gene expression of injury marker Lcn2 and Sox9 in kidneys of male Zfp24^PT-/-^ mice was significantly higher compared to controls; females showed no significant changes. Data are mean ± S.D. Statistical significance determined by two-way ANOVA with Tukey’s post hoc test; ns, not significant; *** p < 0.001.

We further examined SOX9 expression after IRI. In male *Zfp24*^PT–/–^ mice, *Sox9* upregulation was significantly impaired, reinforcing ZFP24’s role in male SOX9 regulation (**Fig. 3g**). In females, no differences in SOX9 expression were observed (**Fig. 3g–h**). These results suggest that the ZFP24–SOX9 axis protects male proximal tubules during AKI but is dispensable in females.

### Blunted upregulation of SOX9 target VGF in female kidneys following IRI

The complete set of SOX9 downstream targets in proximal tubular cells under injury conditions has yet to be fully characterized. In a previous study,^48^ we identified the nerve growth factor–inducible gene *Vgf* (non-acronymic; distinct from VEGF) as a SOX9-dependent, stress-responsive gene that is upregulated during ischemic, nephrotoxic, and rhabdomyolysis-associated renal injury. VGF encodes a precursor polypeptide that is cleaved into bioactive peptides involved in regulating neuronal activity, cell survival, progenitor proliferation, and energy homeostasis.^48^ Similar to SOX9, VGF is upregulated and contributes to a protective response during AKI in male mice.

To determine whether reduced SOX9 induction also affects its downstream target *Vgf*, we performed gene-expression analysis in male and female mice subjected to bilateral IRI. We observed robust *Vgf* upregulation in males, whereas induction in females was significantly diminished—over fivefold lower compared to males (**Fig. 4a**). We then generated *Vgf* conditional knockout mice (**Fig. 4b**). Baseline BUN and creatinine levels were unaffected by *Vgf* deletion in either sex (**Supplementary Table 1**). When subjected to bilateral IRI, *Vgf*^PT–/–^ mice exhibited exacerbated AKI in males, while females showed no difference (**Fig. 4c–g**). These data demonstrate that both the upstream regulation of SOX9 and its downstream targets are differentially controlled between males and females in response to kidney injury.

**Figure 4.**
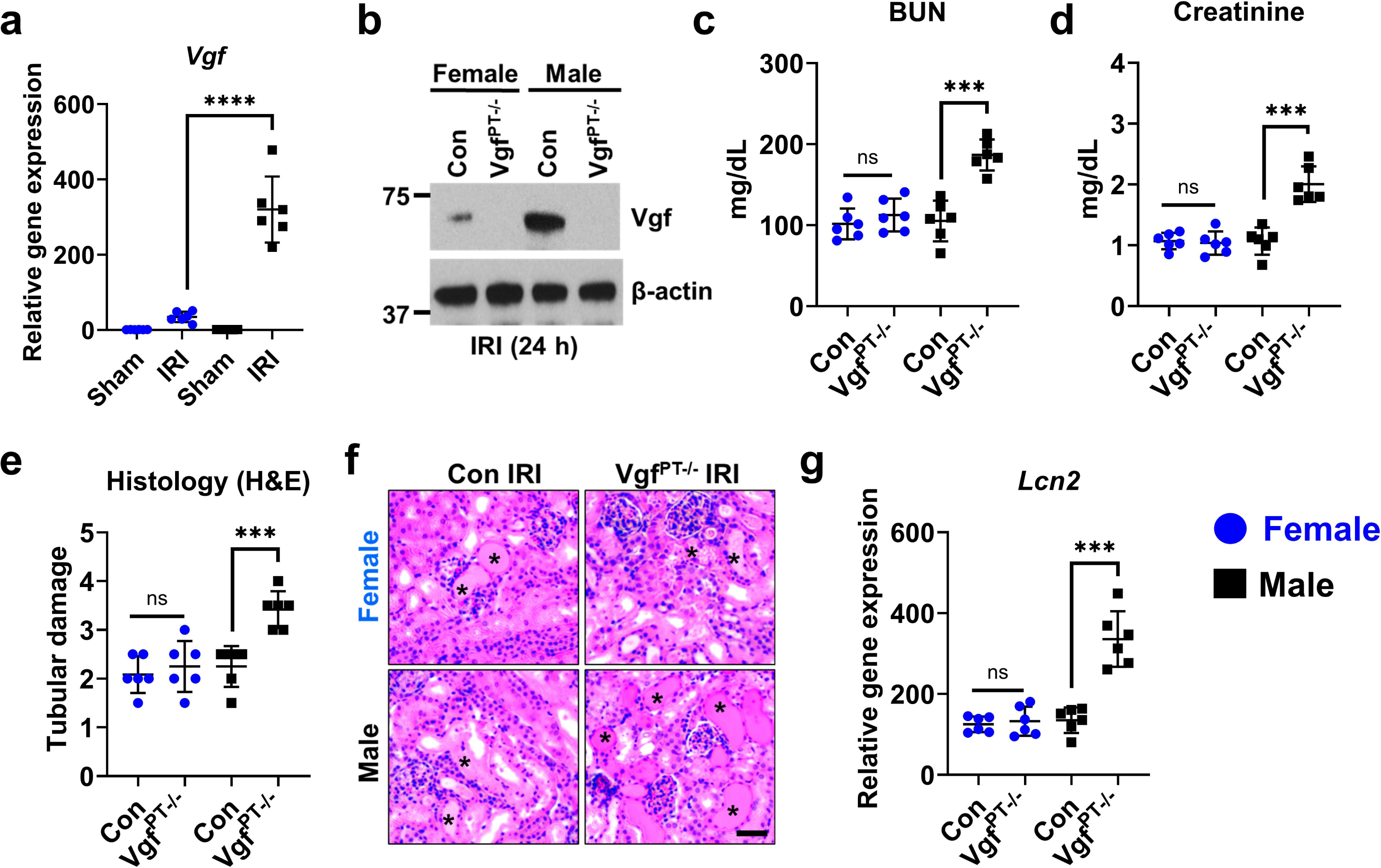
Tubular Vgf deletion protects males but not females from ischemic kidney injury. Tubular epithelial–specific Vgf knockout mice (Vgf^PT-/-^) were generated by crossing Ggt1-Cre mice with Vgf-floxed mice to achieve tubular epithelial–specific Vgf deletion. At 10-12 weeks of age, littermate controls and Vgf^PT-/-^ mice were subjected to bilateral IRI to induce equivalent injury across sexes in the control groups. (**a**) qPCR analysis of Vgf mRNA in kidney lysates from control male and female mice 24 h after bilateral IRI, showing IRI-induced Vgf upregulation predominantly in males. (**b**) Immunoblot of Vgf in kidney lysates from control and Vgf^PT-/-^ male and female mice 24 h after IRI; β-actin is the loading control. (**c–d**) Blood urea nitrogen (BUN) and serum creatinine measurements reveal significantly severe renal injury in Vgf^PT-/-^ male mice compared to controls, with no significant differences observed in females. (**e-f**) Histological tubular injury scores based on H&E staining show reduced tubular damage in Vgf^PT-/-^ males. Representative H&E images depict damaged tubules with an asterisks. Scale bar: 100 µm. (**g**) qPCR analysis of Lcn2 (NGAL) expression confirms elevated injury-marker induction in Vgf^PT-/-^ males but not females. All data are presented as mean ± S.D. with n = 6–8 biologically independent samples per group from three independent experiments. Statistical analysis was performed using two-way ANOVA with Tukey’s multiple-comparison test. Significance is indicated by ***p < 0.001, ****p < 0.0001; ns = not significant.

### SOX4 and SOX11 play protective role in AKI in both male and female mice

Examination of prior bulk RNA-seq datasets ^48^ showed that, along with *Sox9*, *Sox4* and *Sox11* are also upregulated during AKI in male mice. To assess whether these SOX family members are differentially regulated in a sex-specific manner, we first performed qPCR analysis of *Sox4* expression in kidney cortical tissue from male and female mice with matched IRI-induced AKI. Unlike *Sox9*, *Sox4* was upregulated to similar levels in both male and female kidneys (**Fig. 5a**). To determine its functional relevance, we generated conditional *Sox4* knockout mice by crossing *Sox4-floxed* mice with *Ggt1-Cre* mice, confirming SOX4 deletion by Western blot (**Fig. 5b**). These mice showed no renal abnormalities under basal conditions (**Supplementary Table 1**). Following bilateral IRI, *Sox4* deletion led to exacerbated kidney dysfunction in both sexes (**Fig. 5c–g**).

**Figure 5.**
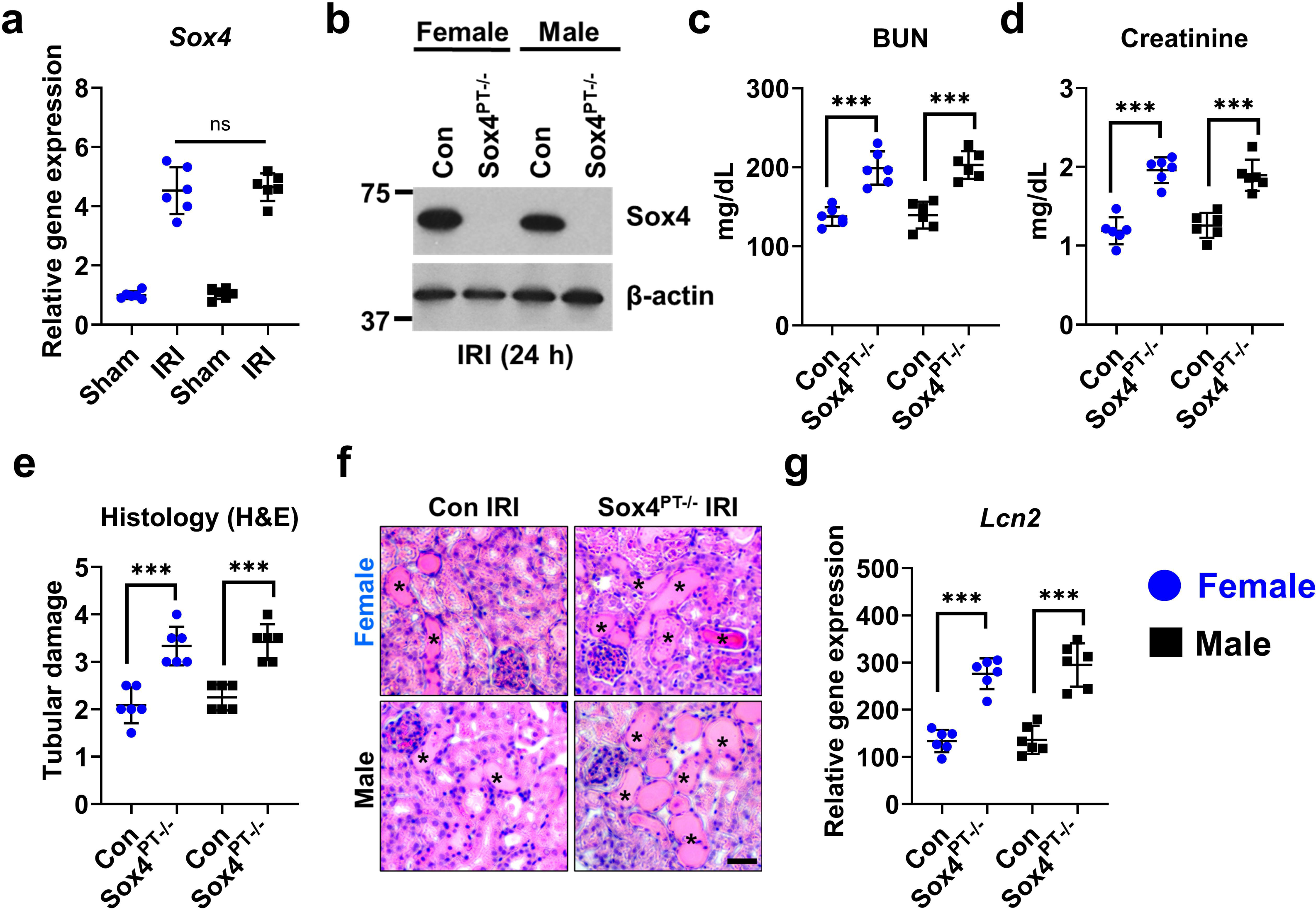
Tubular Sox4 deletion exacerbates ischemic kidney injury in both sexes. Tubular epithelial–specific Sox4 knockout mice (Sox4^PT-/-^) were generated by crossing Ggt1-Cre mice with Sox4-floxed mice to achieve proximal tubular deletion. At 8–10 weeks of age, littermate controls and Sox4^PT–/–^ mice were subjected to bilateral IRI to induce equivalent injury in male and female control mice. (**a**) qPCR analysis of Sox4 expression in kidneys from sham- and IRI-treated control mice (24 h) shows IRI-induced upregulation of Sox4 is not sex-dependent. (**b**) Immunoblot of Sox4 in kidney lysates from control and Sox4^PT-/-^ male and female mice after IRI confirms efficient deletion; β-actin is the loading control. (**c–d**) Blood urea nitrogen (BUN) and serum creatinine measurements reveal significantly worsened kidney function in Sox4^PT-/-^ mice compared to controls in both sexes. (**e-f**) Histological tubular injury scores based on H&E staining confirm increased tubular damage in Sox4^PT-/-^ mice. Representative H&E images depict damaged tubules with an asterisks. Scale bar: 100 µm. (**g**) qPCR analysis of kidney Lcn2 (NGAL) expression shows equivalent injury-marker induction in Sox4^PT-/-^ mice of both sexes compared to controls. All data are presented as mean ± S.D. with n = 5–7 biologically independent samples per group from three independent experiments. Statistical analysis was performed using two-way ANOVA with Tukey’s multiple-comparison test. Significance is indicated by ***p < 0.001; ns = not significant.

Subsequent analysis showed that, similar to *Sox4*, *Sox11* was also upregulated to a comparable extent in males and females (**Supplementary Fig. 6a**). Further studies using proximal tubule–specific *Sox11* conditional knockout mice revealed that *Sox11* also confers protection in both sexes (**Supplementary Fig. 6b–d**). Thus, while *Sox4* and *Sox11* play protective roles in male and female kidneys, *Sox9* appears functionally relevant only in males, highlighting a striking sexual dimorphism in SOX9 regulation and function during AKI.

### Testosterone induces SOX9 upregulation during AKI

To understand the mechanistic basis of sexual dimorphism in *Sox9* regulation during AKI, we considered whether estrogen might inhibit SOX9 upregulation in females or testosterone might promote it in males. To test this, we used gonadectomized mice and optimized clamp times to induce equivalent injury across all groups (**Supplementary Table 2**). Under these matched conditions, AKI severity was comparable in both normal and gonadectomized mice (**Fig. 6a–b, Supplementary Fig. 7**), and NGAL (*Lcn2*) induction was consistent across groups (**Fig. 6c**).

**Figure 6.**
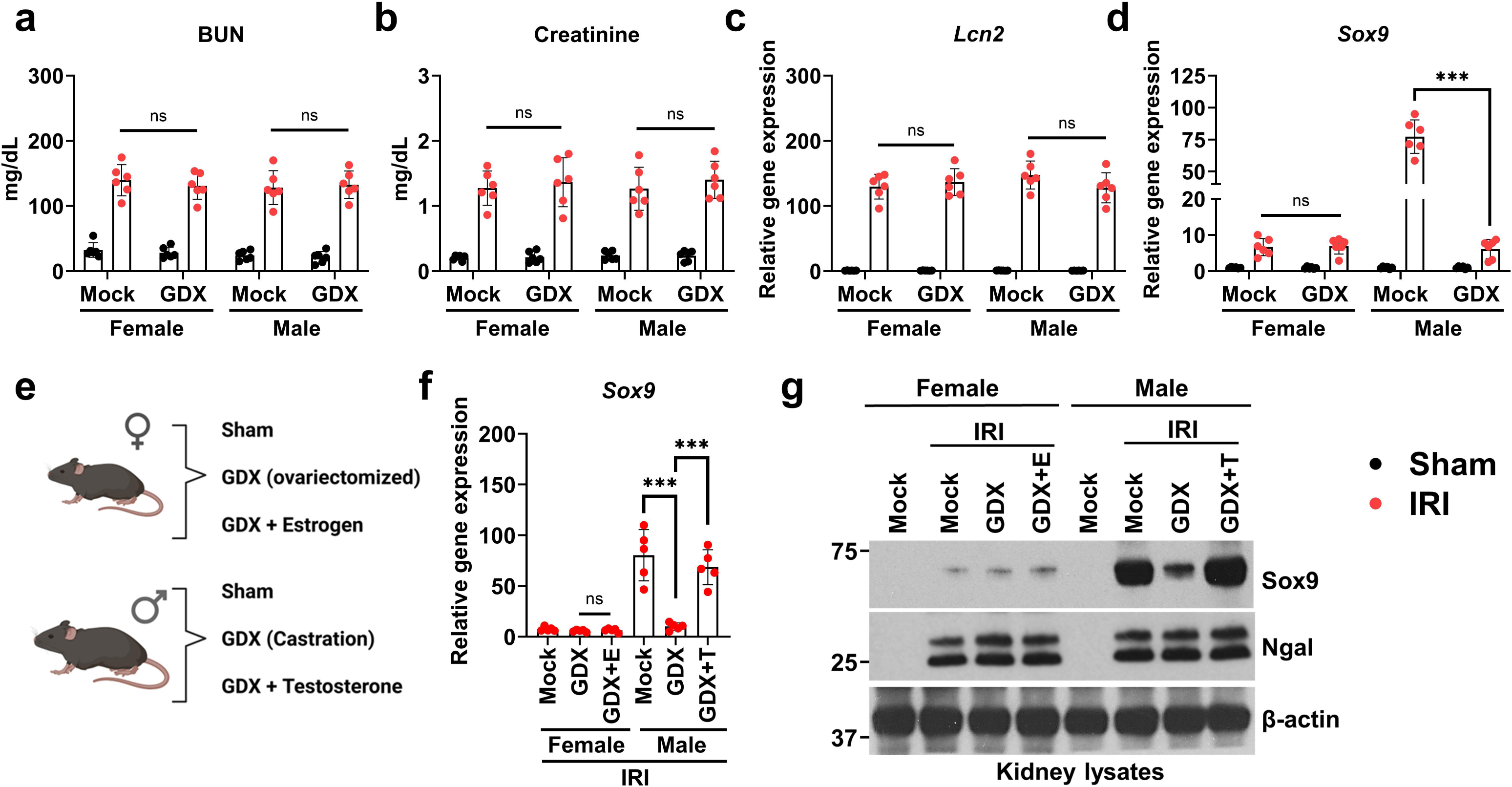
Testosterone drives SOX9 upregulation during ischemic AKI. Age-matched 12-week-old C57BL/6J male and female mice were assigned as Mock (mock surgery) or gonadectomized (GDX) groups, and each cohort was further subjected to either sham surgery or bilateral ischemia–reperfusion injury (IRI) using clamp times optimized to induce equivalent injury in both sexes. At 24 h post-injury: (**a–b**) Blood urea nitrogen (BUN) and serum creatinine confirmed comparable AKI severity across female and male Mock and GDX cohorts.(**c**) Cortical Lcn2 expression showed no sex- or genotype-dependent differences, validating equivalent injury. (**d**) Sox9 induction was markedly diminished in GDX males but unchanged in females. (**e**) Schematic of hormone replacement paradigm: ovariectomized females ± estrogen (E) and castrated males ± testosterone (T). (**f**) Hormone add-back experiments demonstrated that testosterone restored Sox9 induction in GDX males, whereas estrogen had no effect in females. (**g**) Immunoblot analysis of cortical lysates confirmed testosterone-dependent SOX9 induction in males, while NGAL validated equivalent injury across groups; β-actin served as loading control. All data are presented as mean ± S.D. with n = 5–7 biologically independent samples per group from three independent experiments. Statistical analysis was performed using two-way ANOVA with Tukey’s multiple-comparison test. Significance is indicated by ***p < 0.001; ns = not significant.

In male gonadectomized mice, *Sox9* upregulation was markedly reduced, while no change was observed in females (**Fig. 6d**). These data suggest that testosterone drives SOX9 induction in males. To confirm this, we performed hormone add-back experiments in which testosterone or estrogen was administered to gonadectomized males or females, respectively (**Fig. 6e**). Testosterone administration restored *Sox9* upregulation in males, whereas estrogen had no effect in females (**Fig. 6f**). Western blot analysis confirmed that testosterone promotes SOX9 upregulation during AKI (**Fig. 6g**).

### Tubular androgen receptor drives SOX9 upregulation in males

Testosterone exerts its biological effects via the androgen receptor (AR), which is particularly abundant in proximal tubules.^49^ Prior work ^49^ showed that conditional deletion of *Ar* in kidney tubular epithelial cells using *Pax8-Cre* reduces kidney weight in male but not female mice, consistent with a role for AR signaling in the sexually dimorphic increase in kidney size and nephron endowment observed in males. Because *Pax8-Cre* is active during nephrogenesis, those effects likely reflect developmental consequences of AR loss.

To assess the role of AR in injury responses independent of development, we generated a postnatally targeted *Ar* knockout by crossing *Ar-floxed* mice with *Ggt1-Cre* mice, which initiate recombination 7–10 days after birth, after nephrogenesis (**Fig. 7a**). AR deletion (Ar^PT-/-^) did not affect kidney size or baseline renal function in either sex (**Supplementary Table 1**). Following bilateral IRI, male Ar^PT-/-^ mice exhibited reduced tubular injury compared with control males, while females showed no genotype-dependent differences, consistent with the known effects of testosterone on ischemic susceptibility.^20^

**Figure 7.**
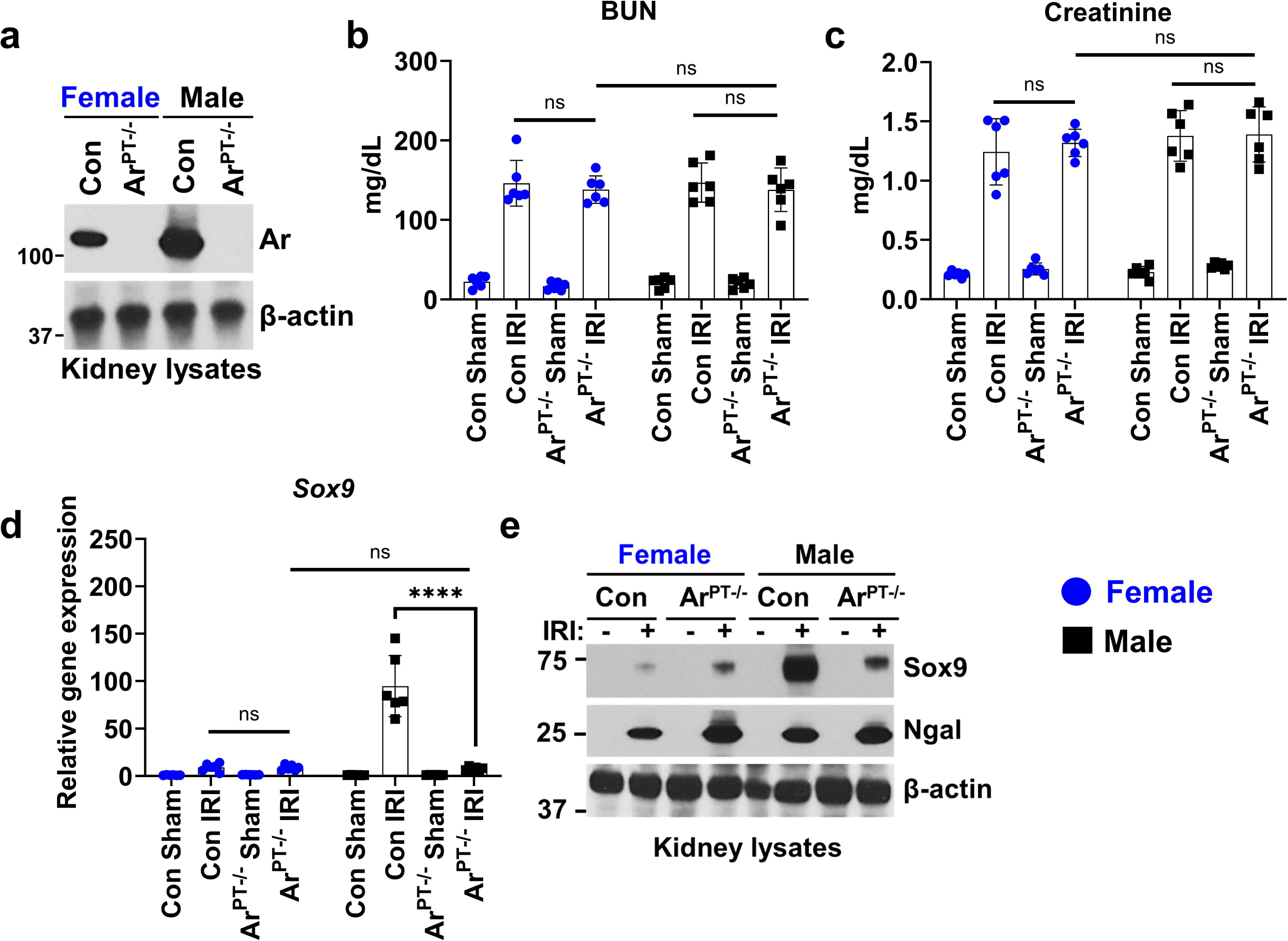
Tubular androgen receptor is required for SOX9 induction in males during ischemic AKI. Tubular epithelial–specific AR knockout mice (Ar^PT-/-^) were generated by crossing Ggt1-Cre mice with Ar-floxed mice to achieve postnatal proximal tubule–specific deletion. At 8–10 weeks of age, littermate controls and Ar^PT-/-^ mice were subjected to bilateral ischemia–reperfusion injury under clamp times optimized to induce equivalent injury across sexes in the control groups. (**a**) Immunoblot validation of AR deletion in kidney cortical lysates from male and female Con and Ar^PT-/-^ mice. (**b–c**) BUN and serum creatinine at 24 h confirmed equivalent AKI severity in Con and Ar^PT-/-^ groups across sexes. (**d**) Sox9 gene expression was robustly induced in male Con mice after IRI but significantly blunted in Ar^PT-/-^ males, with no AR-dependent difference were observed in females. (**e**) Immunoblot analysis of cortical lysates confirmed impaired SOX9 induction in Ar^PT-/-^ males, whereas NGAL expression validated equivalent injury; β-actin served as loading control. All data are presented as mean ± S.D. with n = 5–7 biologically independent samples per group from three independent experiments. Statistical analysis was performed using two-way ANOVA with Tukey’s multiple-comparison test. Significance is indicated by ****p < 0.0001; ns = not significant.

To determine whether AR regulates tubular repair, we adjusted IRI clamp times (**Supplementary Table 2**) to ensure equivalent injury across sexes and genotypes (**Fig. 7b–c**). In control males, *Sox9* expression was robustly induced following injury (**Fig. 7d**). In contrast, male Ar^PT-/-^ mice exhibited significantly blunted *Sox9* upregulation, while females showed no AR-dependent differences. Immunoblot analysis confirmed these results (**Fig. 7e**). These findings demonstrate that AR signaling is required for injury-induced SOX9 activation in a male-specific manner, identifying a direct role for androgen signaling in promoting sex-dependent tubular repair.

## DISCUSSION

This study reveals a fundamental sexual dimorphism in proximal tubular epithelial injury programs. Across ischemic and nephrotoxic models adjusted for equivalent injury severity, male proximal tubules exhibited robust SOX9 induction, whereas female tubules showed markedly attenuated expression. Functionally, *Sox9* deletion exacerbated AKI in males but had no effect in females. These findings challenge the assumption that mechanisms identified in males are directly applicable to females and reinforce sex as a key determinant of tubular stress responses and repair pathways.

SOX9 is minimally expressed in the adult kidney under basal conditions but is strongly induced after injury, where it governs epithelial proliferation, survival, and dedifferentiation. ^21,22^ Previous work in males demonstrated a biphasic role for SOX9, promoting epithelial repair during the acute phase and contributing to fibrosis during maladaptive recovery. ^26^ The present findings add a new dimension to this paradigm by showing that female kidneys mount effective reparative responses without significant SOX9 activation. The blunted upregulation and absence of a phenotype following female-specific *Sox9* deletion indicates that the female kidney utilizes alternative transcriptional programs to maintain epithelial integrity and restore function, underscoring that the molecular circuitry of repair differs fundamentally between sexes.

Mechanistically, this study defines an androgen-dependent regulatory axis that drives SOX9 activation in males. Castration suppressed SOX9 upregulation and testosterone replacement restored it, while ovariectomy had no effect in females. Postnatal PT-specific deletion of the androgen receptor abolished injury-induced SOX9 expression, confirming a direct requirement for testosterone and androgen receptor signaling. This identifies a hormone-driven transcriptional circuit selectively engaged in male PTs during AKI. Upstream, the transcription factor ZFP24 functions as an essential activator of SOX9 in males but not in females. Downstream, the stress-inducible gene *Vgf*, a canonical SOX9 target ^48^, was strongly upregulated and cytoprotective in males but minimally expressed and functionally irrelevant in females. In contrast, SOX4 and SOX11 were induced and required in both sexes, suggesting that these factors provide a general epithelial protective program independent of androgen signaling. Together, these data delineate a testosterone, androgen receptor, ZFP24, SOX9, and VGF axis that orchestrates a male-specific tubular repair response.

Testosterone has complex and dose-dependent effects on kidney function. In experimental models, testosterone increases susceptibility to ischemic AKI, ^20^ in part by promoting oxidative stress, endothelial dysfunction, and inflammatory signaling, whereas low-dose androgen replacement can exert vasoprotective and anti-inflammatory effects. ^50^ In humans, testosterone replacement therapy is associated with a modest but significant increase in AKI risk, ^51^ while androgen deprivation therapy in prostate cancer patients also correlates with higher AKI incidence ^52^. These seemingly contradictory observations suggest that both androgen excess and deficiency can adversely affect renal homeostasis, likely reflecting context-, dose-, and timing-dependent effects on tubular, vascular, and immune compartments. The current study supports a biphasic model in which transient androgen receptor activation during the acute phase promotes adaptive epithelial survival through SOX9 induction, whereas sustained androgen signaling during recovery may favor profibrotic remodeling. Dissecting these phase-specific and cell type-specific effects will be critical for defining safe therapeutic windows for androgen modulation on kidney biology and disease.

Other studies support the concept of sex-specific proximal tubular injury responses. For example, deletion of *Gpx4* induces ferroptosis and severe AKI in males but not in females, ^53^ consistent with greater female resistance to oxidative and metabolic stress. These findings align with our observations and suggest that females engage distinct protective pathways. Interestingly, SOX9 is also upregulated in tubular cells during other disease conditions such as polycystic kidney disease ^54^. Our preliminary data (**Supplementary Fig. 8**) in the *RC/RC* mouse model ^55^ of autosomal dominant polycystic kidney disease (ADPKD) demonstrate that females exhibit significantly lower SOX9 induction than males, indicating that similar mechanisms may operate in cystic disease. More broadly, SOX9 is known to regulate regeneration and progenitor cell activity in several non-renal tissues, including the liver and stem cell compartments. ^56,57^ It will be important to determine whether similar sex-dependent regulation of SOX9 contributes to differential injury responses across organ systems. SOX9 is also a master regulator of sex determination ^58^ and nephrogenesis ^34^, and whether sex differences in developmental SOX9 activity contribute to disparities in nephron endowment and kidney size between males and females remains unknown.

A recent study^59^ from Dr. Andrew P. McMahon’s group showed that androgen receptor signaling drives sexually dimorphic gene expression in proximal tubules under basal, non-injury conditions. Building on this, our work identifies SOX9 as an androgen-responsive effector that becomes selectively induced during acute kidney injury, establishing a context-dependent pathway that governs male-specific tubular injury responses. Future studies are required to identify which other genes are differentially regulated in an androgen-receptor dependent manner under injury conditions.

ZFP24 is activated through dephosphorylated in male PTs during AKI, enabling its binding to the Sox9 promoter and driving Sox9 upregulation.^60^ In contrast, here we have found that Sox9 upregulation is markedly blunted in females and occurs independently of ZFP24, indicating a distinct regulatory mechanism. Future mechanistic studies are required to understand how ZFP24 dephosphorylation in males is coupled to androgen receptor signaling and identify phosphatases that mediate its activation. While we have identified a robust sex-specific difference in SOX9 regulation in proximal tubules, future work should define the shared and divergent pathways that drive stress responses in males and females to advance mechanistic understanding and guide therapeutic targeting.

In conclusion, this study identifies a testosterone, androgen receptor, ZFP24, and SOX9 signaling axis as a male specific tubular repair pathway, whereas females rely on distinct SOX9 independent mechanisms. One limitation of this study is that our analyses were restricted to the acute phase of injury and did not assess later fibrotic stages. Future studies will be needed to determine whether SOX9 contributes to fibrotic remodeling in both males and females. Nevertheless, our findings suggest that therapeutics targeting SOX9 activity may benefit males but are unlikely to be effective in females, who depend on alternative, currently unknown pathways.

## DISCLOSURE STATEMENT

All the authors declared no competing interests.

## DATA SHARING STATEMENT

All data supporting this study are included within the manuscript or the Supplementary Information.

## ACKNOWLEDGMENTS

The authors thank The Ohio State University Comprehensive Cancer Center (OSUCCC) for use of the following shared resources: genomics shared resource, microscopy shared resource, and the OSU University Laboratory Animal Resources (ULAR) for housing and care of animals.

## FUNDING INFORMATION

This study was supported by research grants from National Institute of Health (R01DK132230 and R01DK142433: NSP and AB; R56DK140315 to JK). CVA was supported by Chauncey D. Leake Fellowship from Ohio State University. JAS was supported by OSU Presidential Fellowship. IZK was supported by American Society of Nephrology KidneyCure Pre-Doctoral Fellowship.

## AUTHOR CONTRIBUTIONS

CVA, GS, PSP, QX, AS, JAS, VCT, IZK, HJT, SB, AB, NSP, and JK performed experiments, analyzed data, and or assisted with manuscript writing and editing. YS, CCC, DZO, RR, SKM, and BDH provided reagents, performed data analysis, and or assisted with manuscript writing and editing. NSP and JK conceived the study and approved the final version of the manuscript.

## SUPPLEMENTARY INFORMATION

Supplementary Figure 1: Equivalent AKI severity in male and female mice following ischemia– reperfusion injury.

Supplementary Figure 2: Time course of Sox9 induction following ischemia–reperfusion injury.

Supplementary Figure 3: scRNAseq analysis reveals sexually dimorphic Sox9 induction following ischemic AKI.

Supplementary Figure 4: Blunted Sox9 induction in female kidneys across nephrotoxic and septic AKI models.

Supplementary Figure 5: SOX9 deletion worsens cisplatin nephrotoxicity in males but not females.

Supplementary Figure 6: Proximal tubule SOX11 protects against ischemic AKI in both sexes.

Supplementary Figure 7: Histological assessment of tubular injury in control and gonadectomized (GDX) mice following bilateral IRI.

Supplementary Figure 8: Sox9 expression in kidneys from RC/RC autosomal dominant polycystic kidney disease (ADPKD) mice.

Supplementary Table 1: Baseline serum creatinine levels in conditional knockout mice.

Supplementary Table 2: Bilateral IRI clamp time conditions used to induce equivalent injury in both sexes.

## Notes

### Competing Interest Statement

The authors have declared no competing interest.

## REFERENCES

1. Balzer MS, Rohacs T, Susztak K. How Many Cell Types Are in the Kidney and What Do They Do? Annu Rev Physiol. 2022;84:507–531. doi:10.1146/annurev-physiol-052521-121841

2. Hoenig MP, Zeidel ML. Homeostasis, the Milieu Intérieur, and the Wisdom of the Nephron. Clin J Am Soc Nephrol CJASN. 2014;9(7):1272–1281. doi:10.2215/CJN.08860813

3. Bhargava P, Schnellmann RG. Mitochondrial energetics in the kidney. Nat Rev Nephrol. 2017;13(10):629–646. doi:10.1038/nrneph.2017.107

4. Zuk A, Bonventre JV. Acute Kidney Injury. Annu Rev Med. 2016;67:293-307. doi:10.1146/annurev-med-050214-013407

5. Bellomo R, Kellum JA, Ronco C. Acute kidney injury. The Lancet. 2012;380(9843):756-766. doi:10.1016/S0140-6736(11)61454-2

6. Bonventre JV, Yang L. Cellular pathophysiology of ischemic acute kidney injury. J Clin Invest. 2011;121(11):4210–4221. doi:10.1172/JCI45161

7. Linkermann A, Chen G, Dong G, Kunzendorf U, Krautwald S, Dong Z. Regulated Cell Death in AKI. J Am Soc Nephrol JASN. 2014;25(12):2689–2701. doi:10.1681/ASN.2014030262

8. Murugan R, Kellum JA. Acute kidney injury: what’s the prognosis? Nat Rev Nephrol. 2011;7(4):209–217. doi:10.1038/nrneph.2011.13

9. Chawla LS, Eggers PW, Star RA, Kimmel PL. Acute Kidney Injury and Chronic Kidney Disease as Interconnected Syndromes. N Engl J Med. 2014;371(1):58–66. doi:10.1056/NEJMra1214243

10. Sex and gender disparities in the epidemiology and outcomes of chronic kidney disease | Nature Reviews Nephrology. Accessed July 1, 2025. https://www.nature.com/articles/nrneph.2017.181

11. Chesnaye NC, Carrero JJ, Hecking M, Jager KJ. Differences in the epidemiology, management and outcomes of kidney disease in men and women. Nat Rev Nephrol. 2024;20(1):7–20. doi:10.1038/s41581-023-00784-z

12. Curtis LM. Sex and Gender Differences in AKI. Kidney360. 2023;5(1):160-167. doi:10.34067/KID.0000000000000321

13. Soranno DE, Awdishu L, Bagshaw SM, et al. The role of sex and gender in acute kidney injury-consensus statements from the 33rd Acute Disease Quality Initiative. Kidney Int. 2025;107(4):606–616. doi:10.1016/j.kint.2025.01.008

14. Kumar S, Molitoris BA. Renal Endothelial Injury and Microvascular Dysfunction in Acute Kidney Injury. Semin Nephrol. 2015;35(1):96–107. doi:10.1016/j.semnephrol.2015.01.010

15. Basile DP. The endothelial cell in ischemic acute kidney injury: implications for acute and chronic function. Kidney Int. 2007;72(2):151–156. doi:10.1038/sj.ki.5002312

16. Tiwari R, Sharma R, Rajendran G, et al. Postischemic inactivation of HIF prolyl hydroxylases in endothelium promotes maladaptive kidney repair by inducing glycolysis. J Clin Invest. 135(3):e176207. doi:10.1172/JCI176207

17. Ramesh G, Reeves WB. TNF-α mediates chemokine and cytokine expression and renal injury in cisplatin nephrotoxicity. J Clin Invest. 2002;110(6):835–842. doi:10.1172/JCI15606

18. Li L, Okusa MD. Macrophages, Dendritic cells and Kidney Ischemia-Reperfusion Injury. Semin Nephrol. 2010;30(3):268-277. doi:10.1016/j.semnephrol.2010.03.005

19. Ahmed SB. The importance of sex and gender in basic and clinical research. Nat Rev Nephrol. 2024;20(1):2–3. doi:10.1038/s41581-023-00716-x

20. Park KM, Kim JI, Ahn Y, Bonventre AJ, Bonventre JV. Testosterone Is Responsible for Enhanced Susceptibility of Males to Ischemic Renal Injury*. J Biol Chem. 2004;279(50):52282–52292. doi:10.1074/jbc.M407629200

21. Kumar S, Liu J, Pang P, et al. Sox9 Activation Highlights a Cellular Pathway of Renal Repair in the Acutely Injured Mammalian Kidney. Cell Rep. 2015;12(8):1325–1338. doi:10.1016/j.celrep.2015.07.034

22. Kang HM, Huang S, Reidy K, Han SH, Chinga F, Susztak K. Sox9 positive progenitor cells play a key role in renal tubule epithelial regeneration in mice. Cell Rep. 2016;14(4):861–871. doi:10.1016/j.celrep.2015.12.071

23. Humphreys BD. Sox9 flips the switch between regeneration and fibrosis. Kidney Int. 2024;106(5):781–783. doi:10.1016/j.kint.2024.07.002

24. Kim JY, Bai Y, Jayne LA, et al. A kinome-wide screen identifies a CDKL5-SOX9 regulatory axis in epithelial cell death and kidney injury. Nat Commun. 2020;11(1):1924. doi:10.1038/s41467-020-15638-6

25. Kim JY, Bai Y, Jayne LA, Cianciolo RE, Bajwa A, Pabla NS. Involvement of the CDKL5-SOX9 signaling axis in rhabdomyolysis-associated acute kidney injury. Am J Physiol - Ren Physiol. 2020;319(5):F920–F929. doi:10.1152/ajprenal.00429.2020

26. Aggarwal S, Wang Z, Fernandez Pacheco DR, et al. SOX9 switch links regeneration to fibrosis at the single-cell level in mammalian kidneys. Science. 2024;383(6685):eadd6371. doi:10.1126/science.add6371

27. Sex determination and the control of Sox9 expression in mammals - Jakob - 2011 - The FEBS Journal - Wiley Online Library. Accessed July 1, 2025. https://febs.onlinelibrary.wiley.com/doi/10.1111/j.1742-4658.2011.08029.x

28. Vining B, Ming Z, Bagheri-Fam S, Harley V. Diverse Regulation but Conserved Function: SOX9 in Vertebrate Sex Determination. Genes. 2021;12(4):486. doi:10.3390/genes12040486

29. Morais da Silva S, Hacker A, Harley V, Goodfellow P, Swain A, Lovell-Badge R. Sox9 expression during gonadal development implies a conserved role for the gene in testis differentiation in mammals and birds. Nat Genet. 1996;14(1):62–68. doi:10.1038/ng0996-62

30. Sekido R, Lovell-Badge R. Sex determination involves synergistic action of SRY and SF1 on a specific Sox9 enhancer. Nature. 2008;453(7197):930-934. doi:10.1038/nature06944

31. Coran AG, Cabal L, Siassi B, Rosenkrantz JG. Surgical closure of patent ductus arteriosus in the premature infant with respiratory distress. J Pediatr Surg. 1975;10(3):399–404. doi:10.1016/0022-3468(75)90103-7

32. Li Y, Zheng M, Lau YFC. The Sex-Determining Factors SRY and SOX9 Regulate Similar Target Genes and Promote Testis Cord Formation during Testicular Differentiation. Cell Rep. 2014;8(3):723–733. doi:10.1016/j.celrep.2014.06.055

33. Kashimada K, Koopman P. Sry: the master switch in mammalian sex determination. Development. 2010;137(23):3921–3930. doi:10.1242/dev.048983

34. Reginensi A, Clarkson M, Neirijnck Y, et al. SOX9 controls epithelial branching by activating RET effector genes during kidney development. Hum Mol Genet. 2011;20(6):1143–1153. doi:10.1093/hmg/ddq558

35. Jo A, Denduluri S, Zhang B, et al. The versatile functions of Sox9 in development, stem cells, and human diseases. Genes Dis. 2014;1(2):149–161. doi:10.1016/j.gendis.2014.09.004

36. Croft B, Ohnesorg T, Hewitt J, et al. Human sex reversal is caused by duplication or deletion of core enhancers upstream of SOX9. Nat Commun. 2018;9(1):5319. doi:10.1038/s41467-018-07784-9

37. Odum JD, Akhter J, Verma V, et al. Myeloid-specific ferritin light chain deletion does not exacerbate sepsis-associated AKI. Am J Physiol-Ren Physiol. 2024;327(1):F171–F183. doi:10.1152/ajprenal.00043.2024

38. Martinez SA, Karel IZ, Silvaroli JA, et al. Resazurin dye is an *in vivo* sensor of kidney tubular function. Kidney Int. 2025;107(3):508–520. doi:10.1016/j.kint.2024.12.008

39. Silvaroli JA, Bisunke B, Kim JY, et al. Genome-Wide CRISPR Screen Identifies Phospholipid Scramblase 3 as the Biological Target of Mitoprotective Drug SS-31. J Am Soc Nephrol JASN. 2024;35(6):681–695. doi:10.1681/ASN.0000000000000338

40. Kim JY, Jayne LA, Bai Y, et al. Ribociclib mitigates cisplatin-associated kidney injury through retinoblastoma-1 dependent mechanisms. Biochem Pharmacol. 2020;177:113939. doi:10.1016/j.bcp.2020.113939

41. Silvaroli JA, Martinez GV, Vanichapol T, et al. Role of the CDKL1-SOX11 signaling axis in acute kidney injury. Am J Physiol-Ren Physiol. 2024;327(3):F426–F434. doi:10.1152/ajprenal.00147.2024

42. Bai Y, Kim JY, Bisunke B, et al. Kidney toxicity of the BRAF-kinase inhibitor vemurafenib is driven by off-target ferrochelatase inhibition. Kidney Int. 2021;100(6):1214–1226. doi:10.1016/j.kint.2021.08.022

43. Kim J, Kil IS, Seok YM, et al. Orchiectomy attenuates post-ischemic oxidative stress and ischemia/reperfusion injury in mice. A role for manganese superoxide dismutase. J Biol Chem. 2006;281(29):20349–20356. doi:10.1074/jbc.M512740200

44. Dixon EE, Wu H, Muto Y, Wilson PC, Humphreys BD. Spatially Resolved Transcriptomic Analysis of Acute Kidney Injury in a Female Murine Model. J Am Soc Nephrol JASN. 2022;33(2):279–289. doi:10.1681/ASN.2021081150

45. Kirita Y, Wu H, Uchimura K, Wilson PC, Humphreys BD. Cell profiling of mouse acute kidney injury reveals conserved cellular responses to injury. Proc Natl Acad Sci. 2020;117(27):15874–15883. doi:10.1073/pnas.2005477117

46. Pabla N, Dong G, Jiang M, et al. Inhibition of PKCδ reduces cisplatin-induced nephrotoxicity without blocking chemotherapeutic efficacy in mouse models of cancer. J Clin Invest. 2011;121(7):2709–2722. doi:10.1172/JCI45586

47. Iwano M, Plieth D, Danoff TM, Xue C, Okada H, Neilson EG. Evidence that fibroblasts derive from epithelium during tissue fibrosis. J Clin Invest. 2002;110(3):341–350. doi:10.1172/JCI15518

48. Kim JY, Bai Y, Jayne LA, et al. SOX9 promotes stress-responsive transcription of VGF nerve growth factor inducible gene in renal tubular epithelial cells. J Biol Chem. 2020;295(48):16328–16341. doi:10.1074/jbc.RA120.015110

49. Harris AN, Castro RA, Lee HW, Verlander JW, Weiner ID. Role of the renal androgen receptor in sex differences in ammonia metabolism. Am J Physiol-Ren Physiol. 2021;321(5):F629–F644. doi:10.1152/ajprenal.00260.2021

50. Patil CN, Wallace K, LaMarca BD, et al. Low-dose testosterone protects against renal ischemia-reperfusion injury by increasing renal IL-10-to-TNF-α ratio and attenuating T-cell infiltration. Am J Physiol Renal Physiol. 2016;311(2):F395–403. doi:10.1152/ajprenal.00454.2015

51. Lincoff AM, Bhasin S, Flevaris P, et al. Cardiovascular Safety of Testosterone-Replacement Therapy. N Engl J Med. 2023;389(2):107–117. doi:10.1056/NEJMoa2215025

52. Lapi F, Azoulay L, Niazi MT, Yin H, Benayoun S, Suissa S. Androgen Deprivation Therapy and Risk of Acute Kidney Injury in Patients With Prostate Cancer. JAMA. 2013;310(3):289–296. doi:10.1001/jama.2013.8638

53. Ide S, Ide K, Abe K, et al. Sex differences in resilience to ferroptosis underlie sexual dimorphism in kidney injury and repair. Cell Rep. 2022;41(6):111610. doi:10.1016/j.celrep.2022.111610

54. Patel MM, Gerakopoulos V, Lettenmaier B, et al. SOX9-dependent fibrosis drives renal function in nephronophthisis. EMBO Mol Med. 2025;17(6):1238–1258. doi:10.1038/s44321-025-00233-3

55. Dwivedi N, Jamadar A, Mathew S, Fields TA, Rao R. Myofibroblast depletion reduces kidney cyst growth and fibrosis in autosomal dominant polycystic kidney disease. Kidney Int. 2023;103(1):144–155. doi:10.1016/j.kint.2022.08.036

56. Kadaja M, Keyes BE, Lin M, et al. SOX9: a stem cell transcriptional regulator of secreted niche signaling factors. Genes Dev. 2014;28(4):328–341. doi:10.1101/gad.233247.113

57. Kawaguchi Y. Sox9 and programming of liver and pancreatic progenitors. J Clin Invest. 2013;123(5):1881–1886. doi:10.1172/JCI66022

58. Gonen N, Futtner CR, Wood S, et al. Sex reversal following deletion of a single distal enhancer of Sox9. Science. 2018;360(6396):1469-1473. doi:10.1126/science.aas9408

59. Xiong L, Liu J, Han SY, et al. Direct androgen receptor control of sexually dimorphic gene expression in the mammalian kidney. Dev Cell. 2023;58(21):2338–2358.e5. doi:10.1016/j.devcel.2023.08.010

60. Kim JY, Silvaroli JA, Vasquez Martinez G, et al. Zinc finger protein 24-dependent transcription factor SOX9 up-regulation protects tubular epithelial cells during acute kidney injury. Kidney Int. 2023;103(6):1093–1104. doi:10.1016/j.kint.2023.02.026

