## Supplementary Data for "Androgens Drive SOX9 Upregulation in Injured Proximal Tubular Cells"

#### Supplementary Figure 1

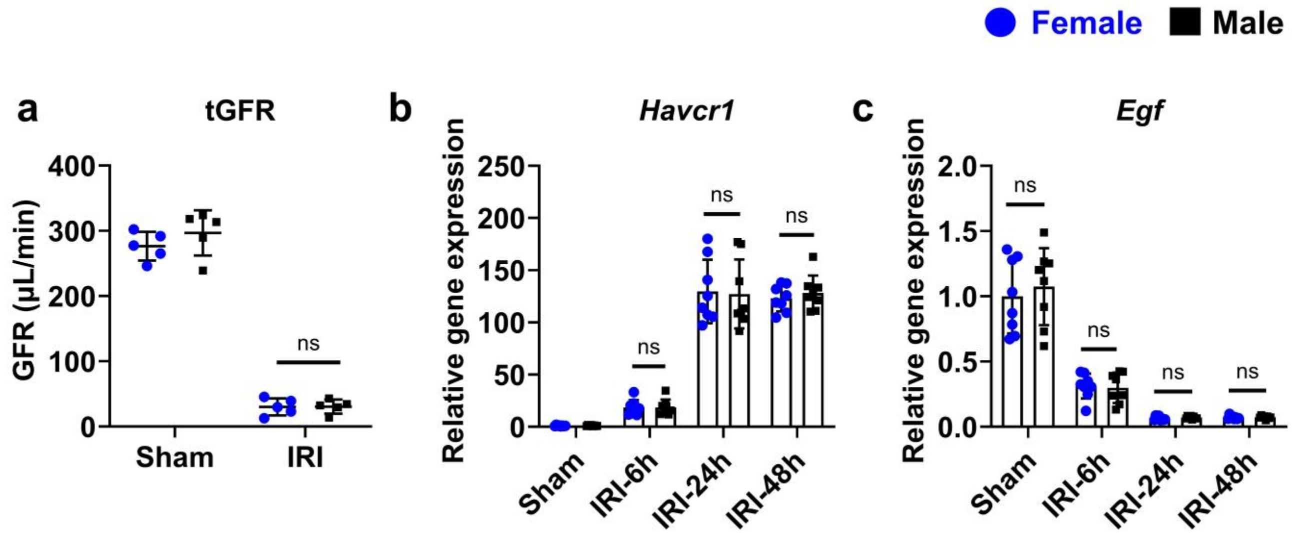


**Supplementary Figure 1. Equivalent AKI severity in male and female mice following ischemia–reperfusion injury.** Age-matched 10–12 week-old C57BL/6J male and female mice were subjected to bilateral ischemia–reperfusion injury (IRI) under clamp times optimized to produce equivalent injury across sexes. At the indicated time points post-IRI: (**a**) Transdermal glomerular filtration rate (tGFR) measurements demonstrated comparable reduction in renal function in males and females. (**b**) Cortical expression of the tubular injury marker Havcr1 (KIM-1) was induced to a similar extent in both sexes at 6 h, 24 h, and 48 h post-IRI. (**c**) Cortical expression of Egf was similarly downregulated in both sexes following IRI. All data are presented as mean ± S.D. with n = 5–7 biologically independent samples per group from three independent experiments. Statistical analysis was performed using two-way ANOVA with Tukey's multiple-comparison test. Significance is indicated by ns = not significant.

#### Supplementary Figure 2

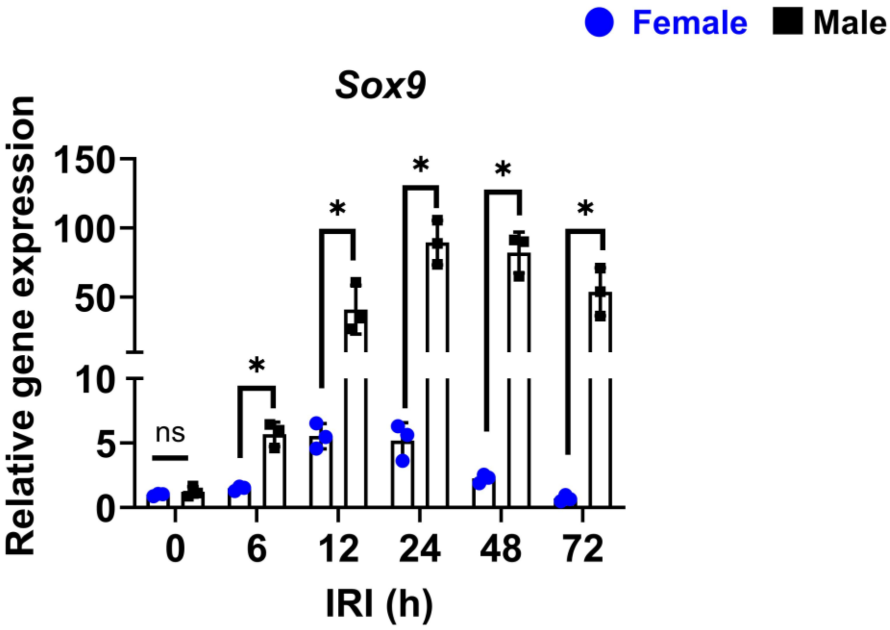


**Supplementary Figure 2. Time course of Sox9 induction following ischemia–reperfusion injury.** Age-matched 10–12 week-old C57BL/6J male and female mice were subjected to bilateral ischemia–reperfusion injury (IRI) under clamp times optimized to induce equivalent injury across sexes. Cortical Sox9 expression was quantified by qPCR at 0, 6, 12, 24, 48, and 72 h post-IRI. Male mice exhibited robust and sustained Sox9 induction from 6–72 h, whereas females displayed consistently blunted responses at all time points. All data are presented as mean ± S.D. with n = 5–7 biologically independent samples per group from three independent experiments. Statistical analysis was performed using two-way ANOVA with Tukey's multiple-comparison test. Significance is indicated by *$p < 0.05$; ns = not significant.

#### Supplementary Figure 3

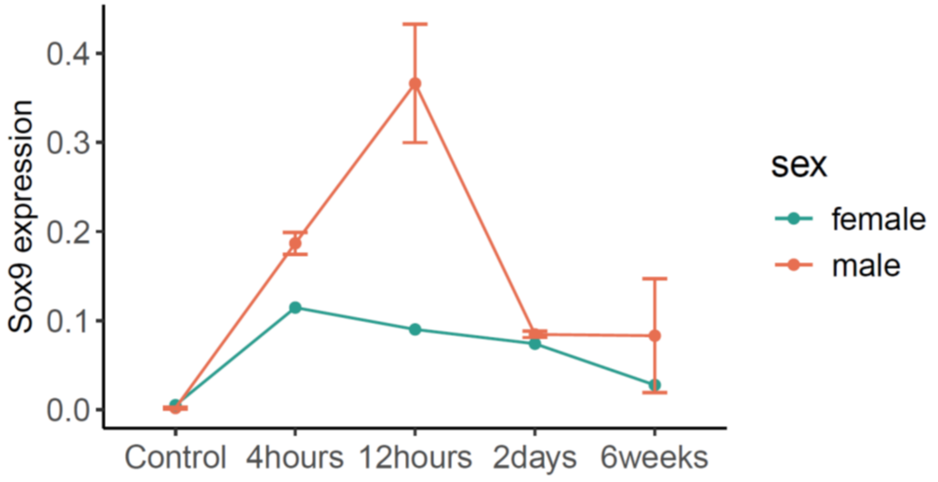


**Supplementary Figure 3. scRNAseq analysis reveals sexually dimorphic Sox9 induction following ischemic AKI.** scRNAseq datasets were reanalyzed to examine Sox9 expression in proximal tubular cells of male and female mice following ischemia–reperfusion injury (IRI). Expression was quantified at control, 4 h, 12 h, 2 days, and 6 weeks post-IRI. Male data were obtained from PNAS, 2020 (PMID 32571916), while female data were derived from JASN, 2023 (PMID 34853151). Male kidneys exhibited a robust and transient peak of Sox9 expression at 12 h, whereas females displayed markedly attenuated induction across all time points. All data are presented as mean ± S.E.M. from integrated single-cell datasets.

#### Supplementary Figure 4

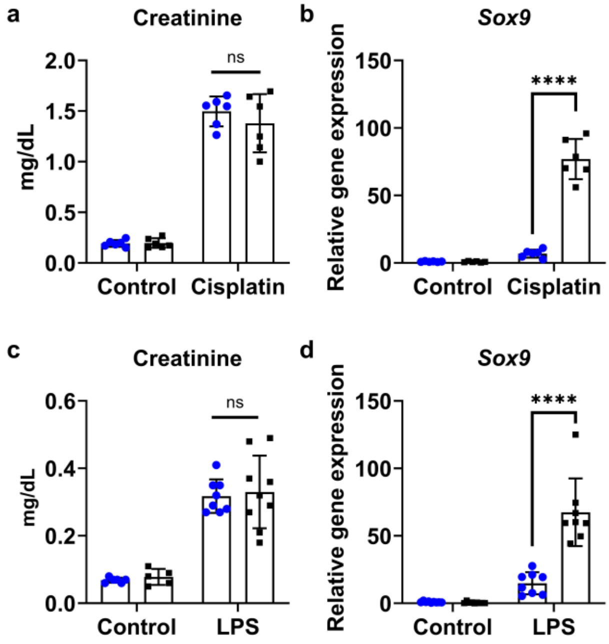


**Supplementary Figure 4. Blunted Sox9 induction in female kidneys across nephrotoxic and septic AKI models.** Age-matched 10–12 week-old C57BL/6J male and female mice were subjected to nephrotoxic or septic AKI under injury conditions optimized to produce comparable renal dysfunction across sexes. (**a–b**) In cisplatin nephrotoxicity, mice received cisplatin by intraperitoneal injection at 30 mg/kg in males and 15 mg/kg in females, and kidneys were harvested at 72 h. Serum creatinine levels were similarly elevated in both sexes, whereas cortical Sox9 expression was robustly induced in males but markedly blunted in females. (**c–d**) In LPS-induced septic AKI, mice were treated with LPS intraperitoneally and kidneys harvested at 24 h. Serum creatinine again rose to comparable levels in males and females, but Sox9 upregulation was significantly greater in males compared with females. All data are presented as mean ± S.D. with n = 5–7 biologically independent samples per group from three independent experiments. Statistical analysis was performed using two-way ANOVA with Tukey's multiple-comparison test. Significance is indicated by ****$p < 0.0001$; ns = not significant.

### Supplementary Figure 5

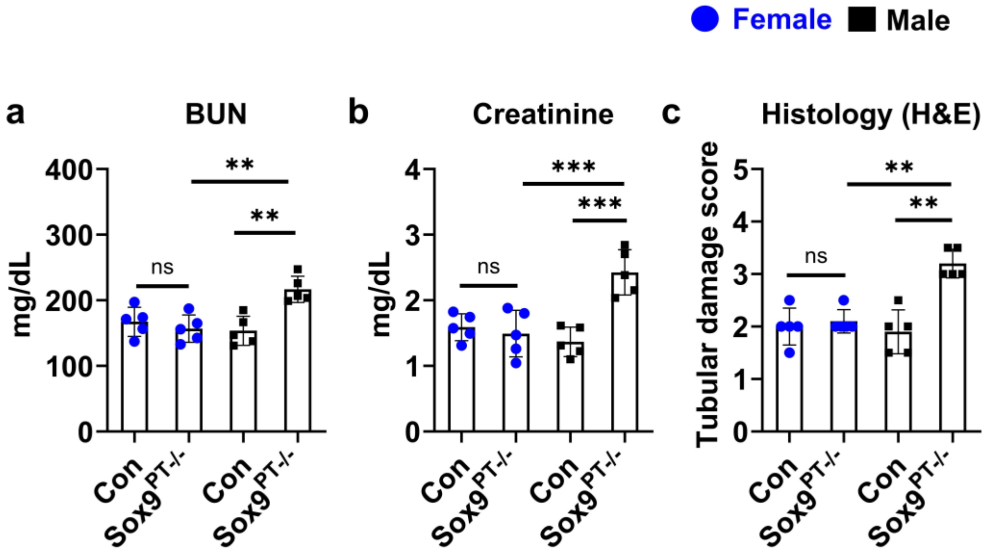


**Supplementary Figure 5. SOX9 deletion worsens cisplatin nephrotoxicity in males but not females.** Tubular epithelial–specific Sox9 knockout mice ($Sox9^{PT-/-}$) were generated by crossing Ggt1-Cre mice with Sox9-floxed mice to achieve proximal tubule–specific deletion. At 10-12 weeks of age, littermate controls and $Sox9^{PT-/-}$ mice were treated with cisplatin (30 mg/kg in males, 15 mg/kg in females; intraperitoneal) and kidneys harvested at 72 h. (**a–b**) Blood urea nitrogen (BUN) and serum creatinine were significantly elevated in male $Sox9^{PT-/-}$ mice compared to male controls, whereas female mice showed no genotype-dependent difference. (**c**) Histological tubular injury scores (H&E) confirmed increased tissue damage in male $Sox9^{PT-/-}$ mice, with no significant difference in females. All data are presented as mean ± S.D. with n = 5–7 biologically independent samples per group from three independent experiments. Statistical analysis was performed using two-way ANOVA with Tukey's multiple-comparison test. Significance is indicated by **p < 0.01, ***p < 0.001; ns = not significant.

#### Supplementary Figure 6

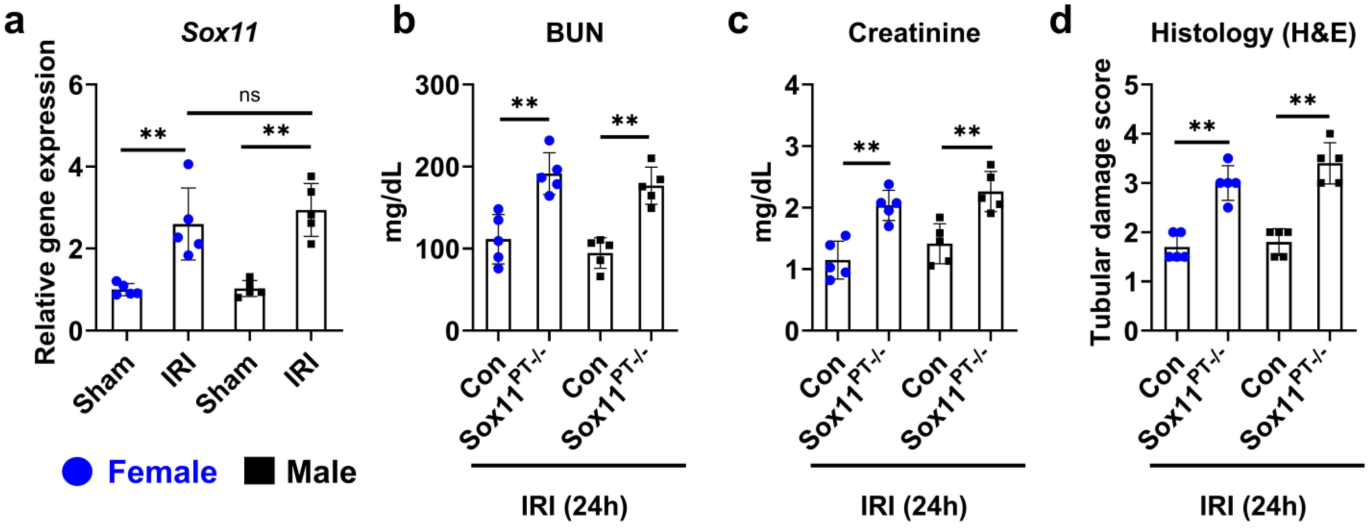


**Supplementary Figure 6. Proximal tubule SOX11 protects against ischemic AKI in both sexes.** Tubular epithelial–specific *Sox11* knockout mice ($Sox11^{PT-/-}$) were generated by crossing Ggt1-Cre mice with Sox11-floxed mice to achieve proximal tubule–specific deletion. At 10–12 weeks of age, littermate controls and $Sox11^{PT-/-}$ mice were subjected to bilateral ischemia–reperfusion injury (IRI) under clamp times optimized to induce equivalent injury across sexes in the control groups. (a) qPCR analysis confirmed Sox11 induction in control mice after IRI, with comparable upregulation in both sexes. (b–c) Blood urea nitrogen (BUN) and serum creatinine at 24 h post-IRI showed significantly worse renal dysfunction in $Sox11^{PT-/-}$ mice compared to controls in both sexes.(d) Histological tubular injury scores (H&E) corroborated increased tissue damage in $Sox11^{PT-/-}$ mice irrespective of sex. All data are presented as mean ± S.D. with n = 5–7 biologically independent samples per group from three independent experiments. Statistical analysis was performed using two-way ANOVA with Tukey's multiple-comparison test. Significance is indicated by **p < 0.01; ns = not significant.

#### Supplementary Figure 7

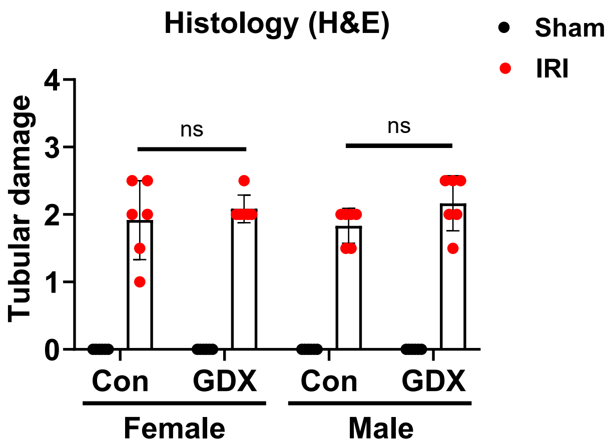


**Supplementary Figure 7. Histological assessment of tubular injury in control and gonadectomized (GDX) mice following bilateral IRI.** Age-matched 12-week-old male and female C57BL/6J mice were subjected to sham surgery (control, Con) or gonadectomy (GDX) prior to sham or IRI procedures. At 24 h post-injury, kidney tissues were collected, stained with hematoxylin and eosin (H&E), and tubular damage was scored based on epithelial flattening, cast formation, tubular dilation, and loss of nuclei. Histological injury scores confirmed equivalent kidney injury across Con and GDX groups following IRI. Data are presented as mean ± S.D. with n = 5–6 biologically independent samples per group from three independent experiments. Statistical analysis was performed using two-way ANOVA with Tukey's multiple-comparison test; $^{**}p < 0.01$, $^{***}p < 0.001$; ns = not significant.

#### Supplementary Figure 8

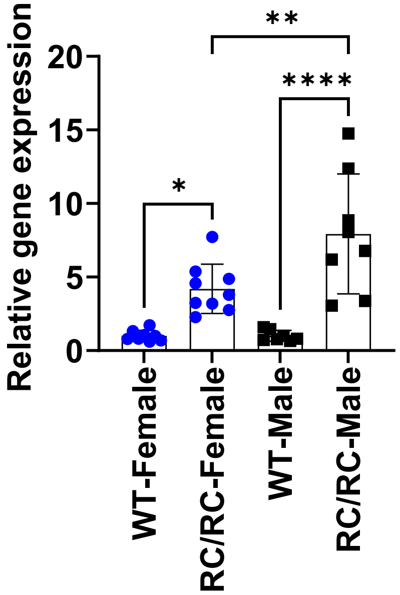


**Supplementary Figure 8. *Sox9* expression in kidneys from RC/RC autosomal dominant polycystic kidney disease (ADPKD) mice.** Quantitative RT-PCR analysis was performed on whole-kidney lysates from wild-type (WT) and RC/RC mice at 10 weeks of age. Both male and female RC/RC mice demonstrated increased *Sox9* expression compared with WT littermates, with significantly higher induction observed in RC/RC males. Data are presented as mean ± S.D. with n = 6–8 biologically independent samples per group. Statistical analysis was performed using one-way ANOVA with Tukey's multiple-comparison test; *p < 0.05, **p < 0.01, ****p < 0.0001.

#### Supplementary Table 1

| | Female | | Male | |
|---|---|---|---|---|
| | **Con** | **Sox9$^{PT-/-}$** | **Con** | **Sox9$^{PT-/-}$** |
| Mean (n=5) | 0.139 | 0.135 | 0.141 | 0.132 |
| SD | 0.026 | 0.024 | 0.033 | 0.042 |
| | **Con** | **Zfp24$^{PT-/-}$** | **Con** | **Zfp24$^{PT-/-}$** |
| Mean (n=5) | 0.158 | 0.141 | 0.150 | 0.162 |
| SD | 0.014 | 0.023 | 0.042 | 0.039 |
| | **Con** | **Vgf$^{PT-/-}$** | **Con** | **Vgf$^{PT-/-}$** |
| Mean (n=5) | 0.128 | 0.134 | 0.139 | 0.143 |
| SD | 0.022 | 0.015 | 0.019 | 0.009 |
| | **Con** | **Sox4$^{PT-/-}$** | **Con** | **Sox4$^{PT-/-}$** |
| Mean (n=5) | 0.141 | 0.154 | 0.163 | 0.134 |
| SD | 0.039 | 0.028 | 0.067 | 0.020 |
| | **Con** | **Ar$^{PT-/-}$** | **Con** | **Ar$^{PT-/-}$** |
| Mean (n=5) | 0.122 | 0.136 | 0.132 | 0.142 |
| SD | 0.022 | 0.029 | 0.035 | 0.030 |

**Supplementary Table 1: Baseline serum creatinine levels in conditional knockout mice.** Serum was collected from 10-12 week old littermate control or conditional knockout mice followed by measurement of serum creatinine levels using the enzymatic method. No significant differences were seen between control and gene knockout mice in both sexes.

#### Supplementary Table 2

| Group | Background | Female (min) | Male (min) |
|---|---|---|---|
| C57BL/6J (WT) | C57BL/6J | 34 | 28 |
| Con and Sox9$^{PT-/-}$ | Mixed (BALB/cJ and C57BL/6) | 36 | 30 |
| Con and Vgf$^{PT-/-}$ | Mixed (BALB/cJ and C57BL/6) | 36 | 30 |
| Con and Zfp24$^{PT-/-}$ | Mixed (BALB/cJ and C57BL/6) | 36 | 30 |
| Con and Sox4$^{PT-/-}$ | Mixed (BALB/cJ and C57BL/6) | 36 | 30 |
| Con and Ar$^{PT-/-}$ | Mixed (BALB/cJ and C57BL/6) | 36 | 30 |
| Mock | C57BL/6J | 34 | 28 |
| GDX | C57BL/6J | 29 | 34 |
| GDX+Testosterone | C57BL/6J | - | 28 |
| GDX+Estrogen | C57BL/6J | 34 | - |

**Supplementary Table 2: Bilateral IRI clamp times (minutes) used to induce equivalent injury in both sexes.**
